# Visualizing transcription sites in living cells using a genetically encoded probe specific for the elongating form of RNA polymerase II

**DOI:** 10.1101/2021.04.27.441582

**Authors:** Satoshi Uchino, Yuma Ito, Yuko Sato, Tetsuya Handa, Yasuyuki Ohkawa, Makio Tokunaga, Hiroshi Kimura

## Abstract

In eukaryotic nuclei, most genes are transcribed by RNA polymerase II (RNAP2). How RNAP2 transcription is regulated in the nucleus is a key to understanding the genome and cell function. The largest subunit of RNAP2 has a long heptapeptide repeat (Tyr1-Ser2-Pro3-Thr4-Ser5- Pro6-Ser7) at the C-terminal domain and Ser2 is phosphorylated on an elongation form of RNAP2. To detect RNAP2 Ser2 phosphorylation (RNAP2 Ser2ph) in living cells, we developed a genetically encoded modification-specific intracellular antibody (mintbody) probe. The RNAP2 Ser2ph-mintbody probe exhibited numerous foci, possibly representing transcription “factories” in living HeLa cells, and foci were diminished when cells were treated with triptolide to induce RNAP2 degradation and with flavopiridol to inhibit Ser2ph. An in vitro binding assay using phospho-peptides confirmed the Ser2ph-specific binding of the mintbody. These results support the view that mintbody localization represents the sites of RNAP2 Ser2ph in living cells. RNAP2 Ser2ph-mintbody foci were colocalized with proteins associated with elongating RNAP2, such as the CDK12 and Paf1 complex component, compared to factors involved in transcription activation around the transcription start sites, such as CDK9 and BRD4. Tracking analysis revealed that RNAP2 Ser2ph-mintbody foci showed constrained diffusional motion like chromatin, but was more mobile compared to euchromatin domains, suggesting that the elongating RNAP2 complexes are separated from the more confined initiating clusters.

**Summary:** The authors developed a genetically encoded probe to specifically detect the Ser2- phosphorylated, elongating form of RNA Polymerase II in living cells. The motion of Ser2- phosphorylated polymerase foci was more dynamic than chromatin domains, suggesting that the elongating complexes are separated from the more confined initiating clusters.

## Introduction

In eukaryotic nuclei, RNA polymerase II (RNAP2) transcribes most genes, and the regulation of RNAP2-mediated transcription is a key to determining the cell phenotype (Cramer, 2019; Reines, 2020; Vos, 2021). Human RNAP2 is a complex of 12 subunits, and the catalytic largest subunit (RPB1) contains a 52 repeat of heptapeptide (Tyr1-Ser2-Pro3-Thr4-Ser5-Pro6-Ser7) at the C-terminal domain (CTD). The site-specific Ser phosphorylation at the CTD is associated with the state of RNAP2 (Eick and Geyer, 2013; Schüller et al., 2016; Zaborowska et al., 2016; Harlen and Churchman, 2017). Unphosphorylated forms of RNAP2 are assembled into preinitiation complexes at the gene promoters, and during the initiation of transcription, Ser5 at the CTD becomes phosphorylated by the cyclin-dependent kinase 7 (CDK7) in the TFIIH complex. RNAP2 Ser5 phosphorylation (Ser5ph) is enriched at the transcription start sites (Komarnitsky et al., 2000) and remains during elongation to facilitate co-transcriptional splicing in mammalian cells (Nojima et al., 2018). After synthesizing a short stretch of RNA, RNAP2 is paused by negative elongation factor and 5,6-dichloro-1-beta-D-ribofuranosylbenzimidazole sensitivity-inducing factor (Yamaguchi et al., 1999; Cramer, 2019; Reines et al., 2020). RNAP2 then undergoes elongation in association with Ser2 phosphorylation (Ser2ph) by positive transcription elongation factor b (P-TEFb), which is a protein complex of CDK9 and Cyclin T (Price, 2000). During elongation, Ser2ph is likely to be maintained by other CDKs, such as CDK12 and CDK13, each complexed with Cyclin K, to facilitate transcription processivity (Fan et al., 2020; Tellier et al., 2020). RNAP2 Ser2ph can mediate the interaction with proteins involved in RNA processing, such as splicing and polyadenylation, as well as epigenetic regulation (Li et al., 2002; Hsin et al., 2012; Gu et al., 2013; Venkat Ramani et al., 2021). RNAP2 becomes dephosphorylated when transcription is terminated. On short genes, such as snRNAs and histones, Ser2 is not phosphorylated (Medlin et al., 2005). Transcription elongation is regulated by many proteins, including the polymerase associated factor 1 (Paf1) complex that was initially identified as a protein complex interacting with RNAP2 in yeast, the depletion of which causes aberrant polyadenylation in mammalian cells (Hou et al., 2019; Francette et al., 2021).

The localization of RNAP2 and its phosphorylated forms in the cell nucleus has been analyzed by light and electron microscopy. RNAP2 Se2ph is closely associated with nascent RNA labeled with Br-UTP, which is consistent with this modification being a mark of elongating form (Iborra et al., 1996; Pombo et al., 1999). From the number of RNAP2 transcription sites (∼8,000) and actively transcribing molecules (∼65,000) in a HeLa cell, it was estimated that one transcription site with ∼80 nm diameter contains eight RNAP2 molecules on average to form transcription “factories” (Jackson et al., 1998; Cook, 1999). In other non-transformed cell types, a smaller number of factories with a larger diameter have been observed (Osborn et al., 2004; Eskiw and Fraser, 2011). In mouse fetal liver erythroblasts, for example, the average diameter of transcription sites is ∼130 nm (Eskiw and Fraser, 2011). The presence of such “factories” can increase the local concentration of the factors involved in transcription, and genes regulated under the same set of transcription factors often share the same factories (Osborn et al., 2004; Xu and Cook, 2008; Schoenfelder et al., 2010; Eskiw and Fraser, 2011; Papantonis et al., 2012).

The dynamic behavior of RNAP2 in living cells has been analyzed using fluorescent protein (FP)-tagged RNAP2 molecules. As more than 100,000 RNAP2 molecules are present in a cell (Kimura et al., 1999; Stasevivch et al., 2014), standard confocal microscopy cannot resolve the small foci (Sugaya et al., 2000; Imada et al., 2021), although the accumulation of FP-tagged RNAP2 can be detected on a transcriptionally activated gene array harboring >100 copies of transcription units (Becker et al., 2002; Janicki et al., 2004). Fluorescence recovery after photobleaching (FRAP) revealed the kinetics of different RNAP2 fractions and the elongation periods (Becker et al., 2002; Kimura et al., 2002; Darzaq et al., 2007; Steurer et al., 2018). High-resolution single-molecule analyses using photo-convertible or -activatable RNAP2 have indicated the transient clustering of RNAP2 during initiation (Cisse et al., 2013; Cho et al., 2016) in association with activators (Boehning et al., 2018) and mediator condensates (Cho et al., 2018). CTD phosphorylation can promote a condensate preference switch from the mediator to splicing factor condensates (Guo et al., 2019). The dynamics of elongating RNAP2 have yet to be documented.

To detect the specifically modified forms of RNAP2 and histones in living cells, fluorescently labeled antigen binding fragments (Fabs) have been used (Hayashi-Takanaka et al., 2011; Stasevich et al., 2014). Using a cell line that harbors a reporter gene array containing glucocorticoid response elements and that expresses glucocorticoid receptor (GR) tagged with GFP, the accumulation kinetics of RNAP2 Ser5ph and Ser2ph on the array upon glucocorticoid stimulation have been revealed (Stasevich et al., 2014). In this system, the target gene array was identifiable by GR-GFP accumulation in a large focus, but Ser5ph and Ser2ph on other single copy genes were not detected. Recently, more detailed kinetic analysis of RNAP2 Ser5 phosphorylation on a single copy gene has been performed, which has also enabled the visualization of the spatiotemporal organization of RNAP2 phosphorylation and mRNA synthesis (Forero-Quintero et al., 2021).

To visualize and track RNAP2 Ser2ph conveniently without protein loading, we developed a genetically encoded live-cell probe, which is a modification-specific intracellular antibody, or mintbody, that consists of the single chain variable fragment (scFv) of the specific antibody and superfolder GFP (sfGFP) (Sato et al., 2013; Sato et al., 2016; Tjalsma et al., 2021). Using high-resolution microscopy, we analyzed the relative localization of RNAP2 Ser2ph-mintbody with proteins involved in RNAP2 phosphorylation, elongation, and transcription activation, such as CDK9, CDK12, a Paf1 complex component Leo1, splicing factor SRRF1/SF2/ASF, bromodomain containing protein 4 (BRD4), and p300 histone acetyltransferase. RNAP2 Ser2ph showed more colocalization with CDK12 and Leo1 than the proteins that facilitate elongation near the transcription start site, such as CDK9, BRD4, and p300. These proteins were more mobile in RNAP2 Ser2ph-enriched regions than elsewhere. RNAP2 Ser2ph foci as such were also more mobile than euchromatin and heterochromatin that were labeled with DNA replication-mediated Cy3-dUTP incorporation. These results suggest that elongating RNAP2 foci are quite mobile compared to the (pre)initiating RNAP2 foci that constrain chromatin motion in living cells.

## Results

### Generation of the RNAP2 Ser2ph-specific mintbody

To generate a mintbody specific for RNAP2 Ser2ph (Fig. 1A), we cloned the coding sequence of antibody heavy and light chains from 18 different hybridoma cells that produce Ser2ph- specific antibodies and transfected scFvs tagged with sfGFP into HeLa cells. Confocal microscopy revealed that a clone, 42B3, was most concentrated in the nucleus among the scFvs, and the others showed weak or little nuclear enrichment (Fig. S1). This suggested that 42B3 scFv was functionally folded and bound to RNAP2 Ser2ph in the nucleus, whereas the majority of scFvs failed to properly fold in the cytoplasm (Wörn et al., 2001; Sato et al., 2013; Sato et al., 2016). In contrast to the sfGFP-tagged version, 42B3 scFv tagged with mCherry (42B3-mCherry) expressed in HeLa cells often exhibited cytoplasmic aggregations and its nuclear enrichment was not as high as that of 42B3-sfGFP (Fig. 1B, top; and S1). Because the folding of scFv is affected by the fusion partner protein or short peptide (Liu et al., 2019), it is likely that sfGFP, but not mCherry, assisted in 42B3 scFv folding. Thus, 42B3 scFv did not appear to have a particularly stable framework, unlike another mintbodies specific for histone H4 Lys20 monomethylation (H4K20me1), whose framework has been used to generate a stable chimeric scFv by implanting the complementary determining regions of a different antibody (Sato et al., 2016; Zhao et al., 2019).

**Figure 1.**
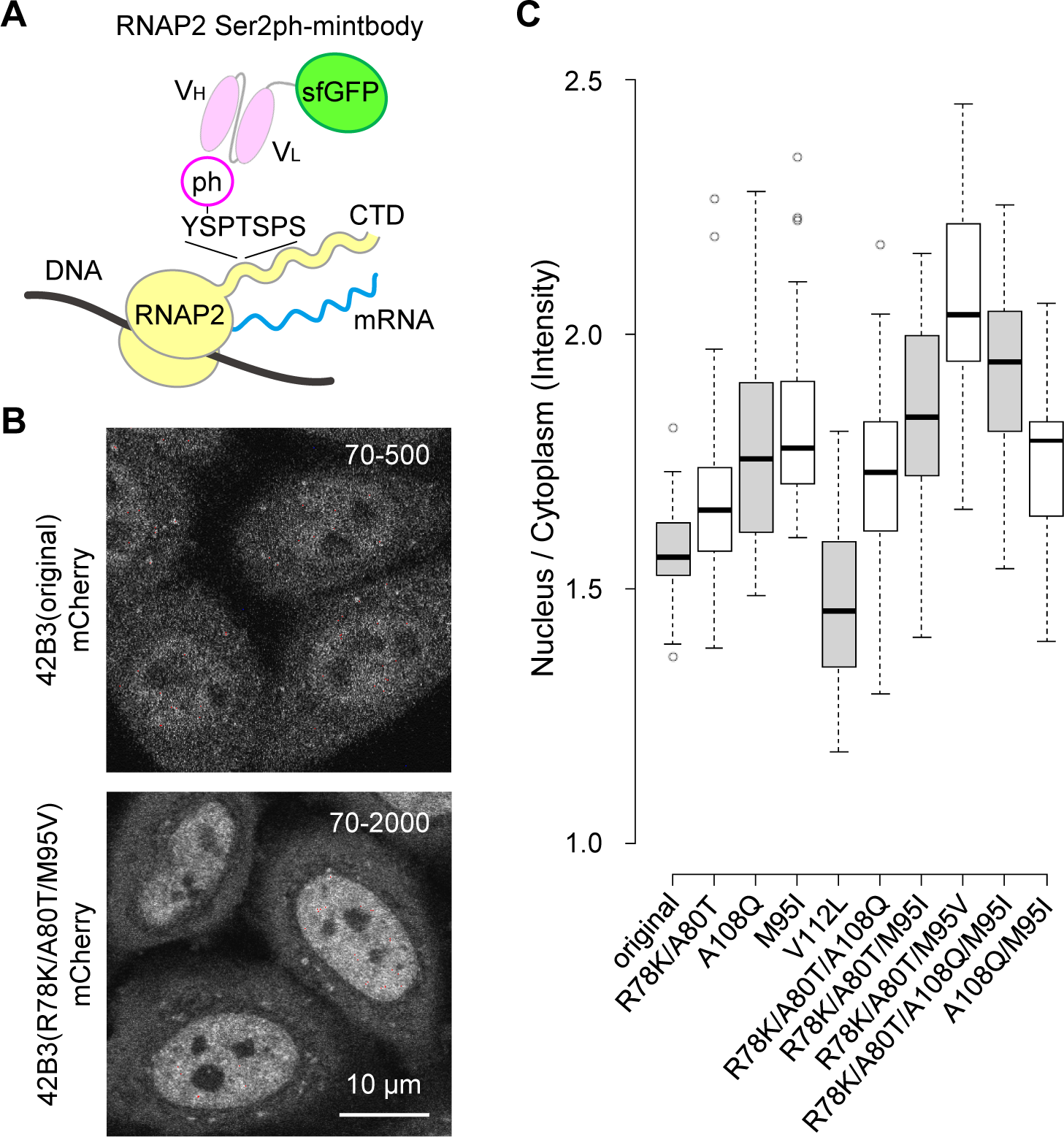
Generation of the RNAP2 Ser2ph-specific mintbody. (A) Schematic diagram of the mintbody that binds to phosphorylated Ser2 on the C-terminal domain of RNA polymerase II. (B) Single confocal sections of 42B3(original)-mCherry and 42B3(R78K/A80T/M95V)-mCherry, which were stably expressed in HeLa cells. Fluorescence images were acquired using a confocal microscope under the same settings, and image contrast was enhanced differently (70-500 for the original 42B3-mCherry and 70-2000 for 42B3(R78K/A80T/M95V)-mCherry) because the intensity of the former was dimmer. Scale bar, 10 µm. (C) Nuclear-to-cytoplasmic intensity ratios of 42B3 and mutants tagged with mCherry. HeLa cells expressing 42B3-mCherry and mutants were established. After confocal images were acquired, the ratios of nuclear-to-cytoplasmic fluorescence intensities were measured (n = 30). In the box plots, center lines show the medians; box limits indicate the 25th and 75th percentiles as determined by R software; whiskers extend 1.5 times the interquartile range from the 25th and 75th percentiles; data points are plotted as open circles.

To improve the folding and/or stability of 42B3 scFv, we introduced amino acid substitutions to the framework region by comparing with the H4K20me1-mintbody (Sato et al., 2016) (Fig. 1C). Five amino acids in 42B3 were replaced with those in the H4K20me1-mintbody individually and in combinations, and the mutant 42B3-mCherry proteins were stably expressed in HeLa cells. By evaluating the nuclear accumulation (i.e., the nucleus/cytoplasmic intensity (N/C) ratio) and the formation of cytoplasmic aggregations, we found that 42B3(R78K/A80T/M95I)-mCherry was the most improved, showing the highest N/C ratio with few cytoplasmic aggregates (Fig. 1C and S1). Structural modeling showed that the R78K/A80T and M95I mutations likely contribute to intramolecular ionic and hydrophobic interactions, respectively. As the hydrophobic cores of scFv are critical for folding and/or stability (Sato et al., 2016; Tjalsma et al., 2021), we examined another M95 mutant with Val substitution and this 42B3(R78K/A80T/M95V)-mCherry exhibited the highest N/C ratio (Fig.1, B and C). Therefore, we used this scFv for subsequent analyses. As described below, 42B3(R78K/A80T/M95V)-sfGFP specifically bound to RNAP2 Ser2ph and so we called this probe RNAP2 Ser2ph-mintbody.

### RNAP2 Ser2ph-mintbody is concentrated in the foci in interphase nuclei in living HeLa cells

We established HeLa cells that stably express RNAP2 Ser2ph-mintbody and observed its distribution using a high-resolution spinning disk confocal microscope (Fig. 2A). RNAP2 Ser2ph-mintbody was concentrated in numerous foci over a diffuse background in the nucleus, suggesting that this probe highlights transcription elongation sites where RNAP2 Ser2ph is present. As one CTD harbors 52 Thy1-Ser2-Pro3-Thr4-Ser5-Pro6-Ser7 repeats and more than a half of Ser2 in the repeats can be phosphorylated (Schüller et al., 2016), a focus may be derived from a single RNAP2 molecule that is bound with multiple RNAP2 Ser2ph-mintbodies. However, it is also possible that a focus may contain multiple polymerases (Iborra et al., 1996). The size of the foci was estimated to be ∼200 nm (Fig. 2B), which is the diffraction limit of confocal microscopy, suggesting that the actual focus size is smaller than 200 nm. The number of foci in a HeLa cell was estimated to be ∼500–5,000 (Fig. 2C), depending on the mintbody brightness. The higher the mintbody fluorescence, the more foci were detected, because the signal-to-background was low in weakly fluorescent cells. The number of foci however appeared to reach a plateau at ∼4,000-5,000, which is close to the number of factories in HeLa cells (Jackson et al., 1998; Pombo et al., 1999).

**Figure 2.**
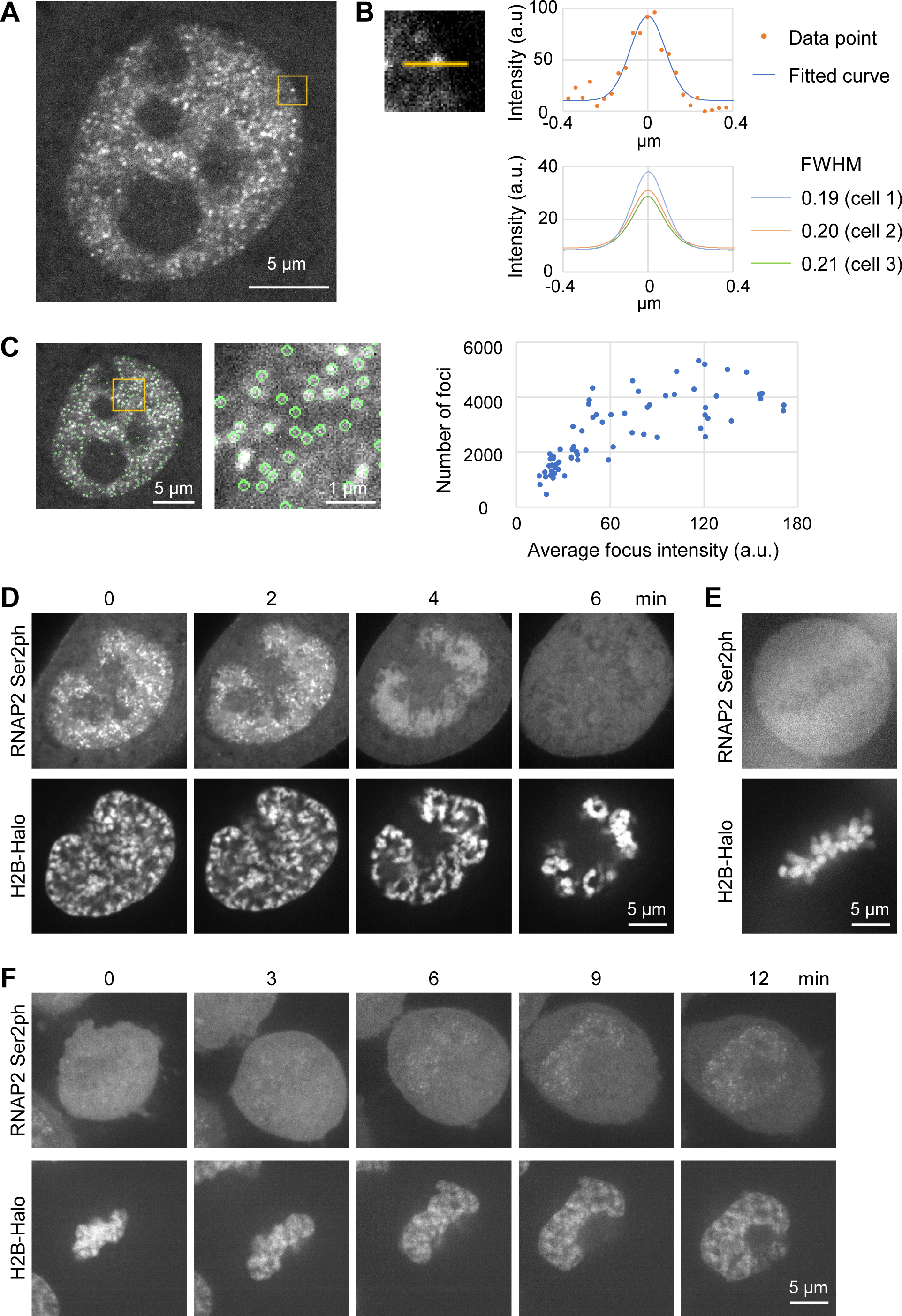
Characterization of RNAP2 Ser2ph-mintbody foci in HeLa cells. (A-C) RNAP2 Ser2ph-mitbody in living HeLa cells. (A) Using a high-resolution spinning disk confocal system, the fluorescence image of RNAP2 Ser2ph-mintbody in a living HeLa cell was acquired. A single section of confocal image is shown. (B) Estimating the size of the foci. Line intensity profiles of single fluorescence foci were plotted in arbitrary units (a.u.). (Top) An example of the focus and its line intensity profile with the fitted curve. (Top left) An example of focus with a 1 µm line in the indicated area in (A). (Top right) The intensity plot of the line and the fitted curve by the Gaussian distribution model. (Bottom) Average fitted curves from 20 spots per cell are shown with an average full width at half maximum (FWHM) for three different cells. (C) Estimating the number of foci. Foci that were identified by a spot detection algorithm are indicated by green circles. A magnified view is shown. The total number of foci were counted by analyzing the z-sections covering the whole nucleus at 0.2 μm intervals. As a single focus can be detected in a maximum of three sections, the total counts were divided by three. The numbers and average intensities (a.u.) of foci in 68 cells are plotted. (D-F) HeLa cells that expressed both RNAP2 Ser2ph-mintbody and Halo-H2B were established. Cells were stained with 50 nM of tetramethylrhodamine-conjugated HaloTag ligand for 30 min before imaging using a high-resolution confocal microscope. Single confocal sections of live-cell images during the (D) prophase to prometaphase, (E) metaphase, and telophase to G1 (F) are shown. (D and F) The elapsed time (min) is indicated. See also Movie 1 and 2 for (D) and (F), respectively. Scale bars, 5 or 1 µm (C; a magnified view).

Most RNAP2 transcription is repressed during mitosis in mammalian cells by the release of the elongation complex (Prescott and Bender, 1962; Parsons and Spencer, 1997; Liang et al., 2015), although a subset of genes can remain to be transcribed (Palozola et al., 2017). By using HeLa cells that express both RNAP2 Ser2ph-mintbody and HaloTag-tagged histone H2B (H2B-Halo), which was labeled with HaloTag tetramethylrhodamine (TMR) ligand to visualize chromatin, we tracked the distribution of RNAP2 Ser2ph-mintbody during mitosis (Fig. 2, D-F; Movie 1 and 2). In cells that started chromosome condensation at the onset of prophase, RNAP2 Ser2ph-mintbody foci were observed around the edge or outside the condensing chromosomes (Fig. 2D, 0 min), which is consistent with the results of a previous study using RNA in situ hybridization to detect nascent transcripts (Liang et al., 2015). The number of foci decreased as the prophase progressed (Fig. 2D, 2 min) and almost disappeared in the late prophase (Fig. 2D, 4 min; Movie 1). After the nuclear membrane broke down, the RNAP2 Ser2ph-mintbody diffused throughout the cytoplasm (Fig. 2D, 6 min). During prometaphase to metaphase, the mintbody appeared to be largely excluded from condensed chromosomes (Fig. 2, E). During the telophase to G1 phase after cytokinesis, RNAP2 Ser2ph- mintbody became concentrated in foci and the number of foci gradually increased in early G1 (Fig. 2F; Movie 2). These mitotic behaviors of RNAP2 Ser2ph-mintbody are consistent with previous observations using fixed cells (Parsons and Spencer, 1997; Liang et al., 2015).

### Mintbody foci depend on RNAP2 Ser2ph in living cells

To investigate whether RNAP2 Ser2ph-mintbody foci in nuclei depend on RNAP2 and its Ser2ph, cells were treated with triptolide, which induces RNAP2 degradation (Wang et al., 2011), and flavopiridol, an inhibitor of the RNAP2 Ser2 kinase P-TEFb (Chao and Price, 2001), respectively. As a control, we also treated cells with the vehicle dimethyl sulfoxide (DMSO). Time-lapse images were collected using a confocal microscope and the total areas of the RNAP2 Ser2ph-mintbody foci were measured (Fig. 3, A and B). Little change in RNAP2 Ser2ph-mintbody foci was observed in cells treated with DMSO for 2 h. In contrast, depletion of RNAP2 with triptolide resulted in the rapid diminishment of nuclear foci. Flavopiridol also induced the diminishment of nuclear foci, but slowly, which is reasonable because elongating RNAP2 remains phosphorylated at Ser2 until it terminates.

**Figure 3.**
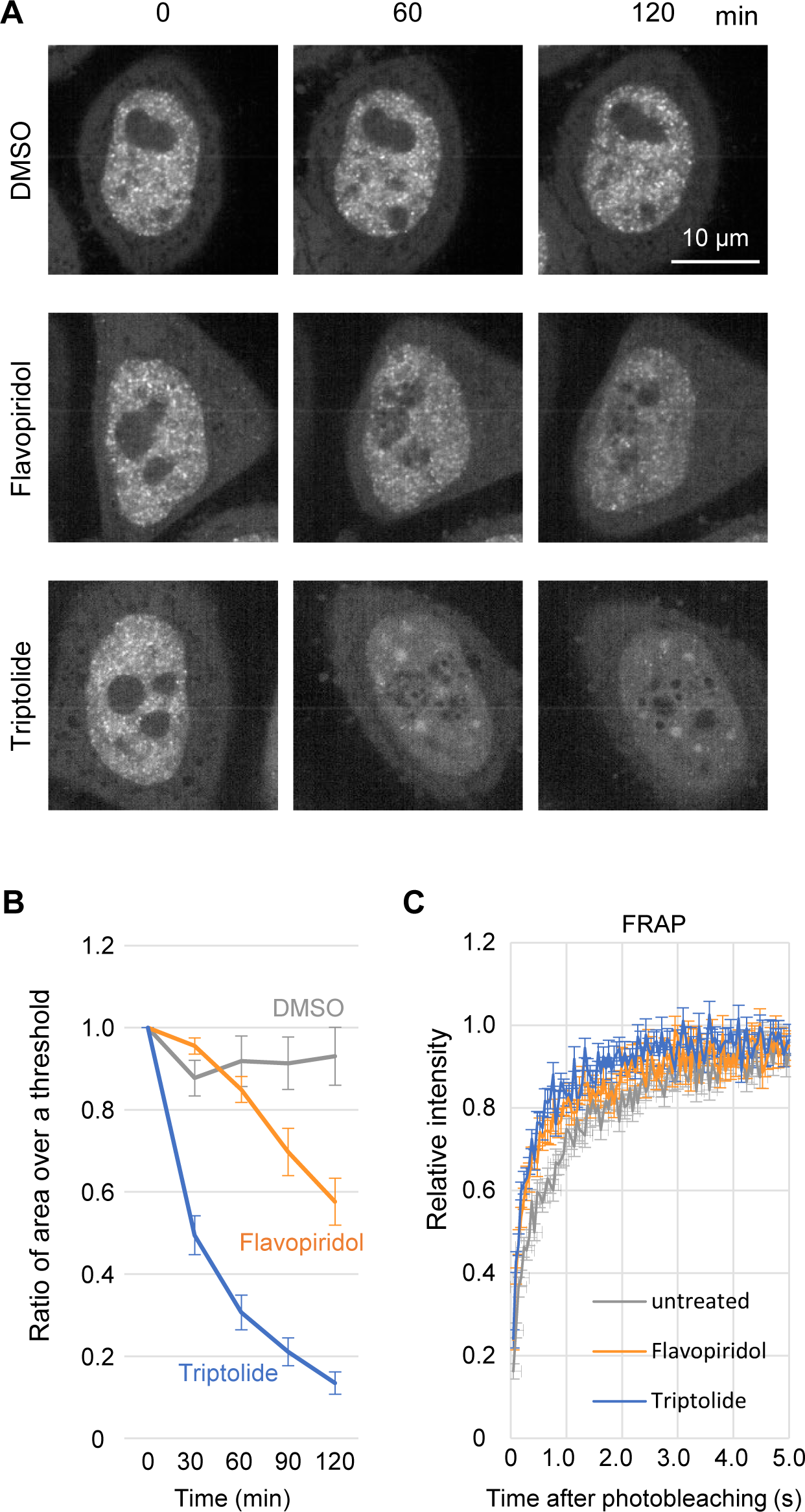
Evaluating the specific binding of the RNAP2 Ser2ph-mintbody to RNAP2 Ser2ph in living cells. (A) and (B) HeLa cells that stably express the RNAP2 Ser2ph-mintbody were treated with 1 µM flavopiridol, 5 µM triptolide, or vehicle (0.1% DMSO) for 2 h. High-resolution confocal images were acquired using a high-resolution confocal system. (A) Representative images at 0, 60, and 120 min. Scale bar; 10 µm. (B) Nuclear foci were selected using an auto-thresholding and the total area in the single nucleus was measured. The relative areas to time 0 were plotted with s.e.m. (n = 10). The area of foci decreased by treatment with flavopiridol and triptolide. (C) Florescence recovery after photobleaching (FRAP). A 2 µm-diameter circle spot in a nucleus was bleached and the fluorescence recovery was measured. RNAP2 Ser2ph- mintbody fluorescence recovered to 80% within 2 s in untreated cells, and the recovery speed increased (to less than 1 s) in cells treated with flavopiridol and triptolide for 2–4 h.

We also used FRAP to examine the kinetics of RNAP2 Ser2ph-mintbody in cells untreated or treated with triptolide and flavopiridol for 2–4 h (Fig. 3C). After bleaching a 2 μm spot in the nucleoplasm, RNAP2 Ser2ph-mintbody signals recovered rapidly to > 80% within 2 s in cells without inhibitors, indicating that this probe binds only transiently to Ser2ph as do the other mintbodies and Fab probes that have been used for tracking posttranslational modifications in living cells without affecting cell division and animal development (Hayashi-Takanaka et al., 2009; Hayashi-Takanaka et al., 2011; Sato et al., 2013; Stasevich et al., 2014; Sato et al., 2016). In triptolide- and flavopiridol-treated cells, the recovery of RNAP2 Ser2ph- mintbody was faster, reaching > 80% within 1 s after bleaching, which was consistent with the decreased level of the target RNAP2 Ser2ph. Thus, kinetic analysis supported the view that RNAP2 Ser2ph-mintbody repeats binding and unbinding to RNAP2 Ser2ph in living cells.

### Biochemical evaluation of RNAP2 Ser2ph-mintbody specificity

To biochemically characterize the specificity of RNAP2 Ser2ph-mintbody, we performed an in vitro enzyme-linked immunosorbent assay (ELISA) using the purified protein. His-tagged RNAP2 Ser2ph-mintbody was expressed in *Escherichia coli* and purified using Ni^2+^ beads. After removal of the His-tag by enterokinase, tag-free RNAP2 Ser2ph-mintbody was further purified (Fig. 4A). ELISA plates were coated with RNAP2 CTD peptides that harbor phosphorylation at different Ser residues and incubated with RNAP2 Ser2ph-mintbody and control IgG antibodies, including the parental 42B3 and previously published Ser2ph (CMA602) and Ser5ph (CMA603) specific antibodies (Stasevich et al., 2014) (Fig. 4B). RNAP2 Ser2ph-mintbody bound to peptides harboring Ser2ph, but not those harboring only Ser5ph and Ser7ph, as did the parental 42B3 IgG and CMA602. These results indicate that RNAP2 Ser2ph-mintbody selectively recognizes Ser2ph at RNAP2 CTD without being particularly affected by Ser5ph and Ser7ph on the same repeat unit. Taken together with cell-based analyses, we concluded that RNAP2 Ser2ph-mintbody can be used as a probe to track RNAP2 Ser2ph in living cells.

**Figure 4.**
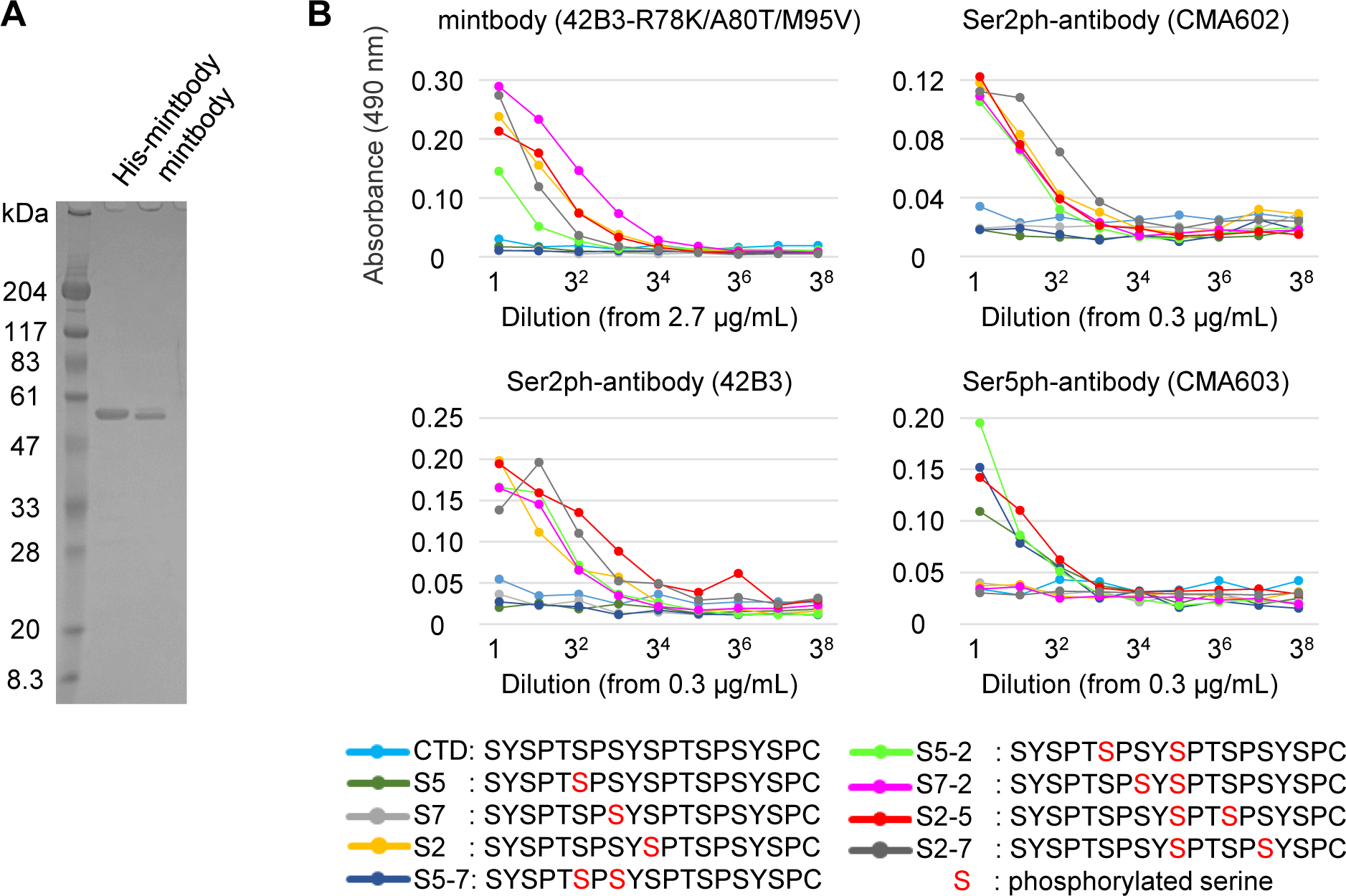
Evaluating the specific binding of RNAP2 Ser2ph-mintbody to phosphopeptides in vitro. RNAP2 Ser2ph-mintbody was expressed in *E. coli* as the His-tag form, purified through an Ni- column, treated with enterokinase to remove Hit-tag, and further purified. (A) Purified proteins were separated on an SDS-polyacrylamide gel and stained with Coomassie Blue. Positions of size standards are indicated on the left. (B) ELISA plates that were coated with synthetic peptides conjugated with BSA, incubated with a dilution series of purified RNAP2 Ser2ph-mintbody, the parental 42B3 antibody (IgG), and control antibodies specific for RNAP2 Ser2ph (CMA602) and RNAP2 Ser5ph (CMA603). After incubations with peroxidase-conjugated anti-GFP (for RNAP2 Ser2ph-mintbody) or anti-mouse IgG (for monoclonal antibodies) and then with o-phenylenediamine, absorbance at 490 nm was measured. RNAP2 Ser2ph-mintbody reacted with peptides containing Ser2ph.

### Localization of transcription-related factors with respect to RNAP2 Ser2ph foci

We compared the localization between RNAP2 Ser2ph-mintbody and proteins that are involved in transcription activation and elongation. An RNAP2 subunit, RPB3, histone H2B, and proliferating cell nuclear antigen (PCNA) were used as positive and negative controls. HaloTag-tagged RPB3 (RPB3-Halo) labeled with TMR showed substantial overlapping with RNAP2 Ser2ph-mintbody (Fig. 5, first column, merged image and line profile). To quantitatively evaluate the colocalization, we employed a cross-correlation function (CCF) analysis, which measures Pearson’s correlations by shifting the image of one fluorescence channel to the x-direction (van Steensel et al., 1996; Fig. 5, bottom row). When foci in two fluorescence images overlap, a peak appears in the middle without shift. The steepness of the peak can also be an indication of a colocalized area. If the distribution of the two images is mutually exclusive, a dip in the middle is expected, and if the distribution is not correlated, either a peak or a dip would not be observed. In the case of RNAP2 Ser2ph-mintbody and RPB3-Halo, a narrow peak in CCF was observed (Fig. 5; first column, bottom), which was consistent with the presence of Ser2ph on the RNAP2 complex that contains RPB3-Halo. The brightest foci of RPB3-Halo without RNAP2 Ser2ph-mintbody (Fig. 5, first column, top, arrowheads) probably represented large RNAP2 condensates, which are often associated with Cajal and histone locus bodies lacking Ser2ph (Xie and Pombo, 2006; Imada et al., 2021). By contrast, H2B- Halo exhibited a complementary pattern to RNAP2 Ser2ph-mintbody (Fig. 5, second column) showing a dip in CCF (Fig. 5, second column, bottom), which agrees with the decondensation of chromatin at transcribed regions. Halo-PCNA exhibited various patterns, depending on the cell cycle stage. Halo-PCNA foci during the mid to late S when heterochromatin was replicated also showed anti-correlation with RNAP2 Ser2ph-mintbody (Fig. S3). Thus, CCF analysis using live imaging data could indicate a relationship between a protein and RNAP2 Ser2ph- enriched transcription foci.

**Figure 5.**
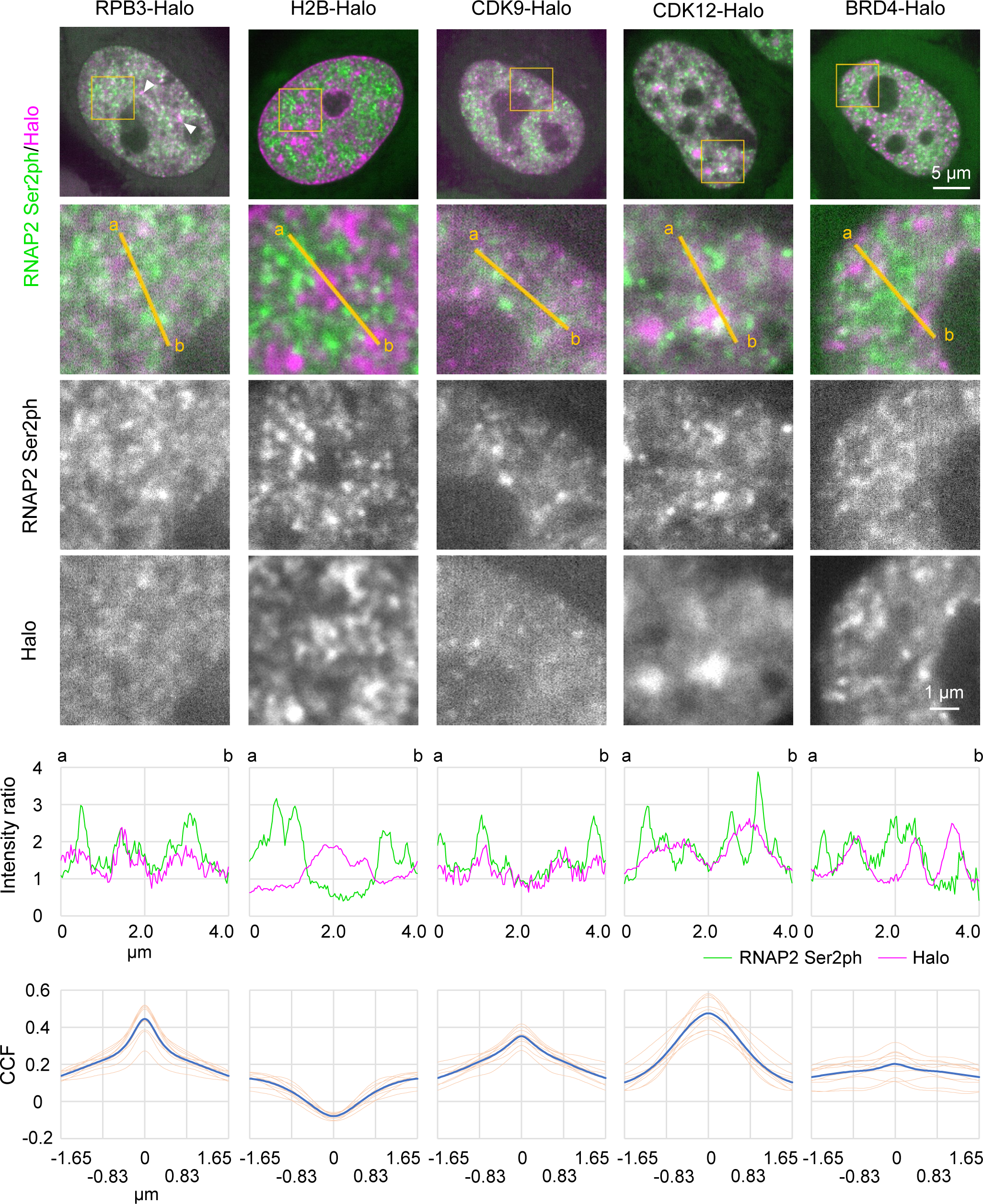
Localization of various proteins with respect to RNAP2 Ser2ph-mintbody. HeLa cells stably expressing RNAP2 Ser2ph-mintbody were transfected with expression vectors for HaloTag- or mRFP-tagged proteins and stained with 50 nM HaloTag TMR ligand for 30 min, before live-cell imaging using a high-resolution confocal system. The averaged images of 10 consecutive frames (551 ms/frame) are shown with magnified views of areas indicated in orange (middle three row). Line intensity profiles of 4 µm lines in the second row are shown (fifth row) by normalizing against the average nuclear intensity to yield relative intensity ratios. The CCFs of 10 cells are shown at the bottom (blue, average; orange, individual cells). See also Fig. S3 for p300, LEO1, SRSF1, and PCNA. Scale bar; 5 µm.

Several kinase complexes phosphorylate RNAP2 Ser2. CDK9 facilitates processive transcription by phosphorylating RNAP2 paused near the transcription start sites (Price, 2000), while CDK12 and CDK13 interact with RNAP2 during elongation (Fan et al., 2020; Tellier et al., 2020). RNAP2 Ser2ph-mintbody foci appeared to be only weakly associated with CDK9-Halo, with a small peak in CCF (Fig. 5, third column). A previous study has also shown infrequent overlapping between CDK9-mCherry and RNAP2 Ser2ph by immunofluorescence (Ghamari et al., 2013). RNAP2 Ser2ph-mintbody foci were rather associated with CDK12-Halo, although the distribution of CDK12-Halo was fuzzier than that of the mintbody, which resulted in a broad CCF peak (Fig. 5, fourth column). CDK12 was also concentrated in splicing factor speckle-like structures (Fig. 5, fourth column, top). A Paf1/RNAP2 complex component, LEO1, showed some overlapping with RNAP2 Ser2ph-mintbody with a CCF peak (Fig. S3). These results suggested that CDK12 and the Paf1 complex are associated, but not always, with RNAP2 Ser2ph. A splicing factor, SRSF1/SF2/ASF1, is recruited to the nascent RNA synthesis sites through the association with RNAP2 CTD phosphorylation (Misteli and Spector, 1999; Gu et al., 2013). RNAP2 Ser2ph-mitbody was often observed at the edge of SRSF1-enriched splicing speckles, which resulted in a broad peak in CCF (Fig. S3).

Histone H3 acetylation is involved in transcription activation through chromatin decondensation and recruiting the activators and bromodomain proteins (Stasevich et al., 2014; Cho et al., 2016; Boehning et al., 2018; Cho et al., 2018). We compared RNAP2 Ser2ph- mintbody localization with an acetyl reader protein, BRD4 (Fig. 5, fifth column), and an acetyltransferase, p300 (Fig. S3), which are associated with enhancers and promoters that harbor H3 Lys27 acetylation. Although the foci between RNAP2 Ser2ph-mintbody and BRD4 or p300 were sometimes closely associated, their overlapping was rare and the correlations were low, which was in good agreement with recent high-resolution analyses showing a separation of BRD4 and enhancers from nascent transcripts (Li et al., 2019; Li et al., 2020).

### Mobility of proteins in RNAP2 Ser2ph-enriched regions

We next analyzed the mobility of single protein molecules inside and outside RNAP2 Ser2ph- mintbody-enriched regions by labeling HaloTag-tagged proteins with a low concentration of the HaloTag TMR ligand using a highly inclined and laminated optical sheet (HILO) microscope (Tokunaga et al., 2008; Lim et al., 2018). As the individual RNAP2 Ser2ph-mintbody molecules bound to the target in less than a few seconds, we did not undertake single-molecule analysis of the mintbody. Instead, we used bulk mintbody fluorescence to define RNAP2 Ser2ph- enriched regions using a laser power much lower than that used in the single-molecule analysis. Under the conditions used, individual RNAP2-mintbody foci were difficult to resolve, thus, mintbody-enriched regions were defined by intensity.

Single HaloTag-tagged proteins were simultaneously imaged with bulk RNAP2 Ser2ph-mintbody at 33.33 ms per frame for 16.67 s (Fig. 6A). The diffusion coefficients (*D*) of the individual molecules during 200 ms were determined by curve fitting (Fig. 6B). The histograms of *D* represent the bimodal distribution of slow and fast fractions (Fig. 6C). As most H2B-Halo molecules were classified into slow fractions (Fig. 6C and S4), this fraction represented the molecules bound to chromatin and/or other large structures and we called this the bound fraction. Relatively large proportions of BRD4 and p300 were also found in the bound fraction compared to RPB3 and CDK9 (Fig. 6C and S4). This probably reflects the small fraction of RNAP2 being the elongating form (Kimura et al., 2002) and the transient interaction of CDK9 with RNAP2.

**Figure 6.**
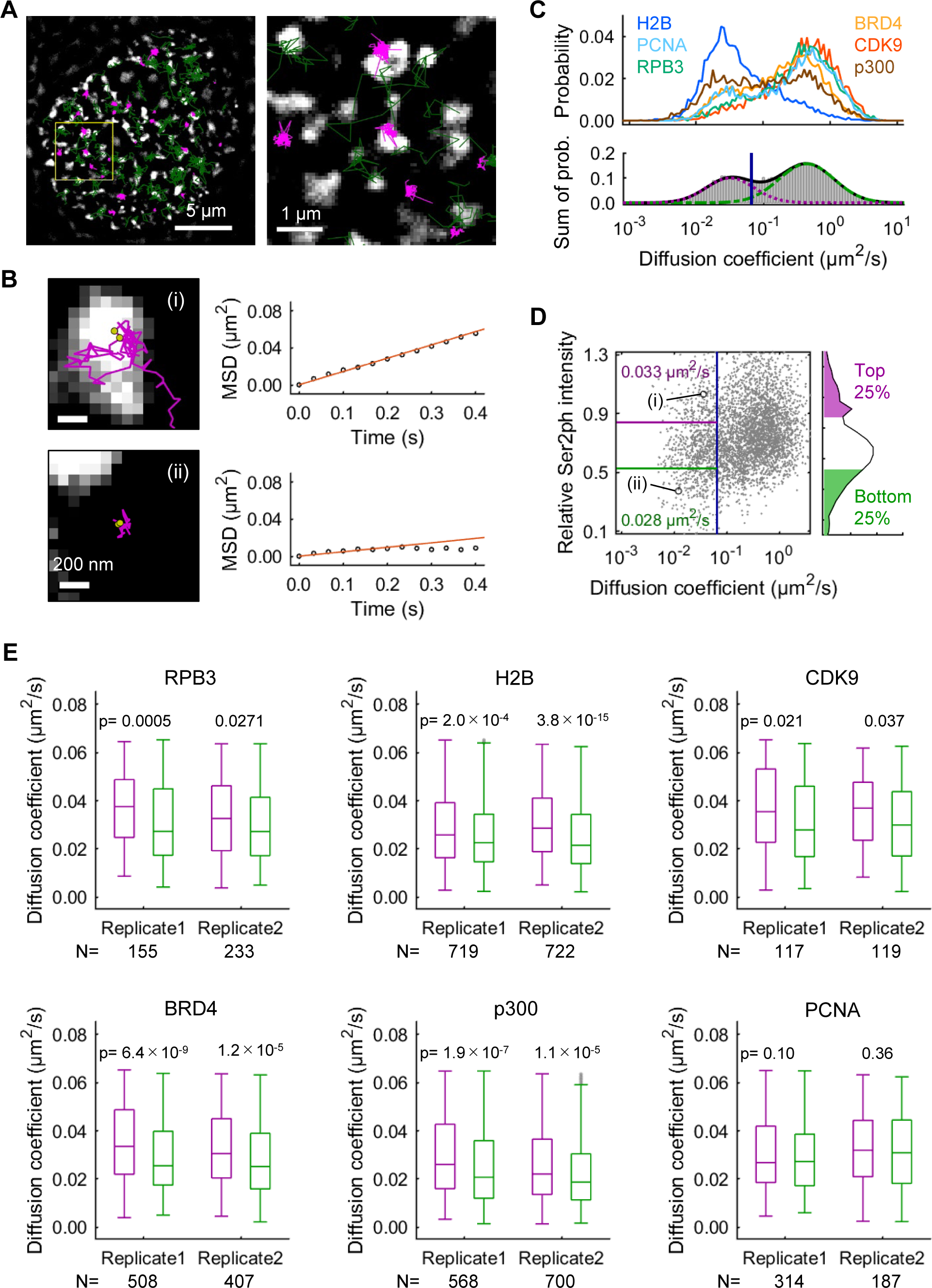
Single-molecule analysis of protein mobility relative to RNAP2 Ser2ph- enriched regions. The mobility of proteins inside and outside RNAP2 Ser2ph foci was quantified using single- molecule trajectories of HaloTag-tagged proteins (RPB3, BRD4, CDK9, p300, H2B, and PCNA) stained with HaloTag TMR ligand recorded at 33.33 ms/frame, which were superimposed upon highpass-filtered RNAP2 Ser2ph-mintbody images in living HeLa cells. (A) Representative single-molecule trajectories of RPB3 superimposed upon RNAP2 Ser2ph- enriched regions (white). Green and magenta lines indicate trajectories of mobile and bound molecules, respectively. Bars, 5 μm (left) and 1 μm (right; a magnified view). (B) Representative trajectories and MSD curves of bound RPB3 molecules inside (i) and outside (ii) RNAP2 Ser2ph-enriched regions. Yellow points represent the tracking points with the top two RNAP2 Ser2ph-mintbody intensities, which were averaged and used as the relative RNAP2 Ser2ph-mintbody intensity (*I*rel_Ser2ph) of the trajectory. The diffusion coefficient *D* was obtained by the linear fitting of MSD (≤ 0.2 s). Bars, 200 nm. (C) The distributions of *D* of the HaloTag-tagged proteins (top) and schematic representation (bottom) showing how to classify the mobile (right) and bound (left) molecules using two- component Gaussian fitting of the distribution of log10(*D*). Fitted lines for two-component Gaussian (solid) and each component (green broken and magenta dotted) are indicated. A blue vertical line indicates the threshold *D*thr by which 97.5% of mobile molecules fall into the mobile fraction. (D) *D* and *I*rel_Ser2ph (left) and the distribution of *I*rel_Ser2ph (right) of the bound molecules (*D* ≤ *D*thr) of RPB3. Each dot on the left represents a single trajectory (molecule). Dots (i) and (ii) correspond to molecules shown in B. Trajectories of the top 25% and the bottom 25% in the *I*rel_Ser2ph distribution (bound) were predominantly located inside and outside RNAP2 Ser2ph- enriched regions, respectively, as shown in B. The average *D*s of bound molecules of the top and bottom 25% *I*rel_Ser2ph are shown in magenta and green, respectively. The blue vertical line indicates *D*thr. The top 25% and bottom 25% fractions in the histogram on the right are colored magenta and green, respectively. (E) *D* of bound HaloTag-tagged proteins, compared between the inside (magenta, top 25% *I*rel_Ser2ph) and outside (green, bottom 25% *I*rel_Ser2ph) of RNAP2 Ser2ph-enriched regions with p values by Mann-Whitney U test. Box plots from two independent replicate experiments are shown. Center lines show the medians; box limits indicate the 25th and 75th percentiles; whiskers extend 1.5 times the interquartile range from the 25th and 75th percentiles, outliers are represented by gray dots. The average *D*s and the corresponding scatter plots are shown in Fig. S4.

We then compared the *D*s of the bound molecules inside and outside the RNAP2 Ser2ph-mintbody-enriched regions based on the intensity of mintbody (Fig. 6, D and E; S4). All proteins, except PCNA, showed higher *D* inside RNAP2 Ser2ph-mintbody-enriched regions compared to that outside, suggesting that the elongating RNAP2 and/or the molecules in the clusters were more mobile than chromatin. To further investigate the dynamics of the transcription sites, we analyzed the mobility of RNAP2 Ser2ph-mitbody foci.

### Mobility of RNAP2 Ser2ph foci

To compare the mobility of RNAP2 Ser2ph-mintbody foci with chromatin domains, we labeled replication domains by DNA replication-mediated Cy3-dUTP incorporation (Manders et al., 1999). HeLa cells expressing RNAP2 Ser2ph-mintbody were loaded with Cy3-dUTP to pulse-label replicated chromatin and were further grown for two days. As a limited amount of Cy3- dUTP was loaded into cells, DNA regions that were replicated just after the loading were labeled and then Cy3 signals exhibited characteristic DNA replication foci depending on the stage in the S-phase (Manders et al., 1999). Once incorporated, Cy3 on DNA persisted after cell divisions, thus, the replication domains could be tracked in living cells. We compared the mobility of RNAP2 Ser2ph-mintbody foci and Cy3-DNA domains that showed euchromatic or heterochromatic patterns by imaging at 551 ms/frame for 27 s using a confocal microscope (Fig. 7). RNAP2 Ser2-mintbody foci exhibited constrained diffusional motion with faster mobility than euchromatin (Fig. 7A) and heterochromatin domains (Fig. 7B). Directional movements of RNAP2 Ser2ph-mintbody foci were hardly observed during this period. Taken together with the single-molecule tracking data (Fig. 6), these results suggest that the elongating RNAP2 complexes and molecules therein are not fixed to a single location in the nucleus but rather are more mobile than chromatin domains.

**Figure 7.**
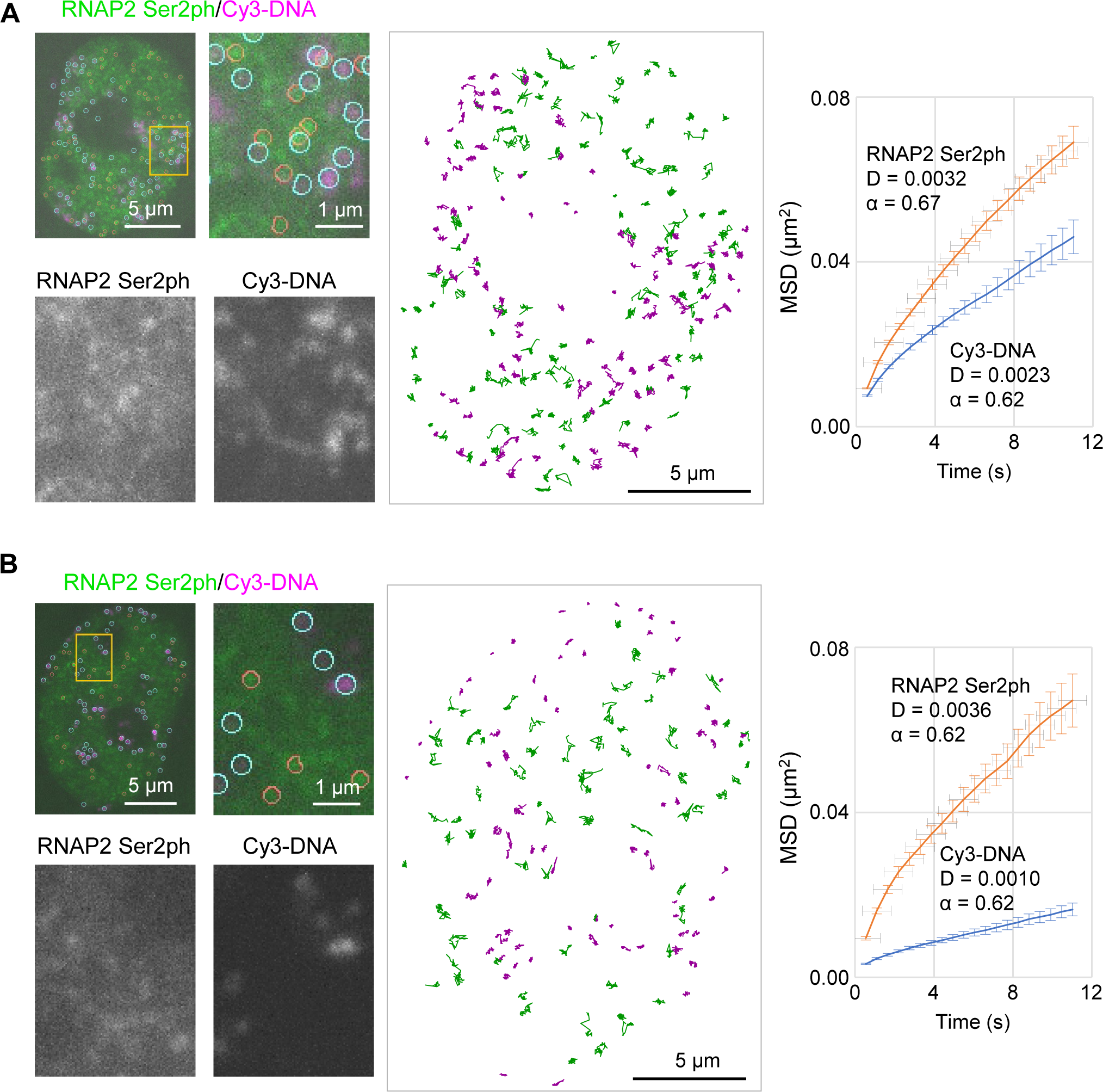
Mobility of RNAP2 Ser2ph foci. HeLa cells stably expressing RNAP2 Ser2ph-mintbody were loaded with Cy3-dUTP to pulse- label chromatin domains that replicated just after loading and cells were further grown for two days. Fluorescent images were acquired at 551 ms/frame using a high-resolution confocal system. RNAP2 Ser2ph-mintbody and Cy3-DNA foci that show euchromatic (A) and heterochromatic (B) patterns were tracked to measure their mobilities. The whole nucleus, magnified views with indications of tracked foci (orange and cyan circles for mintbody and Cy3- DNA, respectively), and traces of individual foci are shown. MSDs are shown (means with s.e.m.) with diffusion coefficients and anomalous factors. Numbers of analyzed foci are 1059 (RNAP2 Ser2ph) and 954 (Cy3) for A, and 379 (RNAP2 Ser2ph) and 716 (Cy3) for B, from nine cells for both A and B. Scale bars, 5 µm (whole nuclei) and 1 µm (magnified views).

## Discussion

### RNAP2 Ser2ph-mintbody probe

In this study, we developed a genetically encoded probe for visualizing Ser2-phosphorylated RNAP2 in living cells based on the specific antibody. The expression of functional scFv depends on the folding and stability of the framework regions (Cattaneo and Biocca, 1999; Ewert et al., 2004; Sato et al., 2016; Tjalsma et al., 2021). The original 42B3 clone showed a moderate level of nuclear enrichment, an indication of the functionality of scFv that targets a nuclear protein or its modification. Introducing several amino acid substitutions improved the nuclear enrichment. Although no universal framework is currently available (Kabayama et al., 2020), some framework sequences are more stable than others and a substitution of the Met residue in a hydrophobic core with a more hydrophobic amino acid, such as Leu and Ile, often increases scFv stability (Tjalsma et al., 2021). In 42B3, Met95 substitution to Val or Ile also improved the nuclear accumulation, implying that such substitutions are a good option to improve the folding and stability of scFv in general. Amino acid substitutions that could affect hydrophilic interactions also improved the 42B3 scFv, but less effectively than the Met substitution in the hydrophobic core. These amino acid substitution results are beneficial for the future development of intracellular antibodies for research and therapeutic purposes (Stocks, 2005; Messer and Butler, 2020).

The resulting RNAP2 Ser2ph-mintbody was concentrated in numerous foci in the nucleus depending on RNAP2 Ser2ph, making it possible to detect and track RNAP2 transcription elongation sites in living cells. Even though the binding residence time of the mintbody was less than a few seconds, keeping the expression level low is important so as to not disturb the turnover of phosphorylation. Therefore, we selected cells that expressed relatively low levels of the mintbody and confirmed the sensitivity to flavopiridol.

### RNAP2 Ser2ph-mintbody foci

Ser2ph is a hallmark of an elongating form of RNAP2 (Eick and Geyer, 2013; Zaborowska et al., 2016; Harlen and Churchman, 2017). Lines of evidence support the view that RNAP2 Ser2ph-mintbody foci in living cells represent the sites of RNA-engaged Ser2-phosphorylated RNAP2. First, the mintbody foci disappeared and reappeared during the prophase to prometaphase and during the telophase to G1, respectively, which is consistent with the substantial repression of RNAP2 transcription during mitosis (Parsons and Spencer, 1997; Liang et al., 2015). Second, the mintbody foci were observed depending on RNAP2 and Ser2ph, demonstrated by inhibitor treatments. The photobleaching assay confirmed the increased mobility of the mintbody due to a loss of the binding target by inhibitors. Third, the binding specificity was validated by an in vitro ELISA using purified protein. Thus, the location of RNAP2 Ser2ph is likely to be detected by the specific mintbody.

As the brightness of single mintbody foci was not at the single-molecule level, multiple mintbody molecules were concentrated in a space less than the ∼200 nm diffraction limit. As Ser2 in the CTD repeat can be highly phosphorylated (Schüller et al., 2016), a single RNAP2 could have ∼50 mintbody binding targets, although it is not known whether the variations at the distal C-terminal repeats, such as Tyr1-Ser2-Pro3-Thr4-Ser5-Pro6-Lys7 and Tyr1-Ser2- Pro3-Thr4-Ser5-Pro6-Thr7, are still recognized. However, it is unlikely that 50 mintbody molecules are housed on a single RNAP2 because of steric hinderance. In addition, if all Ser2ph is occupied by mintbodies, the binding of endogenous proteins can be blocked and dephosphorylation may be inhibited. Although it is still possible to have tens of mintbodies per RNAP2 to exhibit a fluorescence focus, the number of foci in a HeLa cell does not correspond to the number of total active RNAP2 molecules, which is estimated to be tens of thousands (Jackson et al., 1998; Kimura et al., 1999). The maximum number of foci detected (∼5,000) was rather similar to thousands of transcription sites containing multiple transcription units, suggesting that the mintbody foci likely correspond to transcription factories (Iborra et al., 1996; Pombo et al., 1999; Cook, 1999).

### Motion of RNAP2 Ser2ph foci

We observed that protein molecules associated with RNAP2 Ser2ph-rich regions were more mobile than those outside the foci. Similarly, RNAP2 Ser2ph foci were more mobile than chromatin domains. This observation is consistent with the results of a study that showed the higher mobility of transcribed loci (Gu et al., 2018) but not with those of other studies that showed that RNAP2 transcription constrains chromatin motion (Ochiai et al., 2015; Germier et al., 2017; Nagashima et al., 2019). Single-molecule analyses have also revealed that transcription constrains chromatin motion (Nagashima et al., 2019; Shaban et al., 2020). The lower mobility of chromatin is still observed in cells treated with CDK9 inhibitors (Germier et al., 2017; Nagashima et al., 2019), implying that transcription-dependent chromatin constraint is mediated through transcription initiation rather than elongation. As only a small fraction of RNAP2 Ser2ph foci was associated with CDK9 (also demonstrated by Ghamari et al., 2013), most mintbody foci corresponded to RNAP2 Ser2ph that were already elongating on the gene body but not at the transcription start sites. In addition, BRD4 foci were even further apart from RNAP2 Ser2ph or transcripts (Li et al., 2019; Li et al., 2020). Therefore, it is possible that the initiating and elongating RNAP2 clusters are differently organized in space.

Typical reporter genes that are short and highly transcribed (Ochiai et al., 2015; Germier et al., 2017; Forero-Quintero et al., 2021) are likely dominated by the initiating form even though elongating RNAP2 is also present, which may result in confined chromatin motion. The endogenous genes are often periodically transcribed by bursting (Suter et al., 2011, Fukaya et al., 2016; Larsson et al., 2019; Ochiai et al., 2020) and the elongating RNAP2 can be separated from the transcription start sites. In this case, the elongating Ser2- phosphorylated RNAP2 clusters on a single gene by transcription bursting or different genes become free of the confined initiating clusters and exhibit more diffusible motion. The chromatin structure needs to be extensively loosened during ongoing transcription, and the elongating complex on such loosened chromatin fibers can become less constrained. The higher mobility at the single-molecule level may also be associated with the dynamic structure of RNAP2 clusters. It is thus tempting to speculate that clustered RNAP2 molecules that transcribe different genes are pulled through chromatin motions from different angles, resulting in more dynamic mobility than euchromatin domains.

## Materials and methods

### Cells and transfection

HeLa cells were grown in Dulbecco’s modified Eagle’s medium high glucose medium (Nacalai Tesque) containing 10% fetal bovine serum (FBS; Gibco) and 1% L-Glutamine–Penicillin– Streptomycin solution (GPS, Sigma) at 37°C under a 5% CO2 atmosphere. For transfection, Fugene HD (Promega) was used according to the manufacturer’s instructions. Briefly, 2 µg DNA was mixed with 6 µL of Fugene HD in 100 µL of Opti-MEM (Thermo Fisher Scientific) and incubated at room temperature for 30 min, before being added to HeLa cells grown in 35 mm glass bottom dishes (AGC Technology Solutions) to 40-70% confluence. To establish stable cell lines, 1.6 µg PB533-based Piggybac plasmid and 0.4 µg transposase expression vector (Systems Bioscience) were used, and cells were selected in 1 mg/mL G418 (Nacalai Tesque). Cells expressing RNAP2 Ser2ph-mintbody at low levels were collected using a cell sorter (SH800; Sony). Before live-cell imaging, the medium was replaced with FluoroBrite (Thermo Fisher Scientific) containing 10% FBS and 1% GPS.

### Cloning and evaluation of antibody variable fragments encoding 42B3 and its mutants

Mouse hybridoma cell lines expressing antibodies specific to RNAP2 Ser2ph were generated and screened by Mab Institute Inc. as previously described (Kimura et al., 2008). RNA was extracted from the hybridoma cells using TRIzol (Thermo Fisher Scientific) and subjected to RNA-seq to determine the DNA sequence encoding the IgG heavy and light chains, as described previously (Kuniyoshi et al., 2016). The variable regions of heavy (VH) and light (VL) chains of 42B3 were amplified and connected, as described previously (Sato et al., 2018; Tjalsma et al., 2021), using sets of primers specific for VH (VH_s, VH_as), VL (VL_s, VL_as), and the linker (LINK primer1 and LINK primer2), which are listed in Table S1. The resulting scFv fragments were further amplified using the 5’ and 3’ primers (scFv primer_s and scFv primer_as) and cloned into the psfGFP-N1 vector (Addgene #54737, deposited by Michael Davidson and Geoffrey Waldo; Pédelacq et al., 2006) to generate a 42B3-sfGFP expression vector.

Point mutation primers were designed (Table S1) for PCR using 42B3-scFv as a template with Prime STAR (TaKaRa). KOD one PCR Master Mix Blue (Toyobo) was used for the M95V mutation. After PCR reactions, the amplified products were treated with DpnI (0.4 units/μL; 1 h) to digest the template before mixing with competent cells. The nucleotide sequence of the resulting plasmids was verified by Sanger sequencing. 42B3 scFv mutants were then subcloned into a PB533-based vector (Systems Bioscience) by restriction enzyme digestion (EcoRI and NotI) and ligation.

To screen mintbodies and mutants, a point-scan confocal microscope (Olympus; FV- 1000) operated by the built-in software FLUOVIEW ver. 4.2 was used with a UPlanApoN 60× OSC oil immersion lens (NA 1.4) using the following settings: 512 × 512 pixels, pixel dwell time μs, pinhole 100 μm, zoom ×3.0, line averaging ×3, with 543 nm (50%) laser line (Fig. 1B), or zoom ×5.0, line averaging ×4, with 488 nm (3%) laser line (Fig. S1).

To determine the N/C ratio of scFv-mCherry (Fig. 1), the whole cell, nucleus, and cell-free background regions were selected manually and the intensities and areas were measured using ImageJ Fiji 1.52d software (https://imagej.net/Fiji). Net intensities of the whole cell and nucleus were obtained by subtracting background intensity and the cytoplasmic intensity was calculated by dividing the total cytoplasmic intensity ([whole cell intensity] × [whole cell area] – [nuclear intensity] × [nuclear area]) by the cytoplasmic area ([whole cell area] – [nuclear area]). The N/C ratio was obtained by dividing the cytoplasmic intensity by the nuclear intensity.

### Purification of RNAP2 Ser2ph-mintbody

The RNAP2 Ser2ph-mintbody sequence was inserted into pTrc-His (Thermo Fisher Scientific) to harbor the His-tag at the N-terminus. *E. coli* BL21 (DE3) harboring the expression plasmid was grown in 100 mL of YTG (1% tryptone, 0.5% yeast extract, 0.5% NaCl, and 0.2% glucose) for 20 h at 18°C to reach the early stationary phase, diluted to a final volume of 2 L of YTG, and further grown for 18 h at 15°C. Isopropyl-β-D-thiogalactopyranoside was added at a final concentration of 1 mM and *E. coli* culture was further incubated for 12 h at 15°C before collecting cells by centrifugation at 4,000 ×*g* for 10 min at 4°C. After discarding the supernatant, the pellet was frozen at −80°C. The pellet was thawed on ice and then suspended in 20 mL of Buffer L (50 mM Tris-HCl, pH 8.0, 300 mM NaCl, 5% glycerol) containing 1 mg/mL lysozyme (Nacalai Tesque) and 1% proteinase inhibitor cocktail (Nacalai Tesque). Cells were lysed in icy water by sonication using a Sonifier 250 (Branson; duty cycle 30%, output 1.2; 60 s; 30 times with 60 s intervals). The supernatant was collected after centrifugation at 15,000 ×*g* for 15 min at 4°C and applied to a 20 mL open column (Poly-Prep Chromatography Columns; BIO-RAD) containing 500 µL of Ni-NTA agarose (Qiagen) preequilibrated with Buffer L at 4°C. After washing the column twice with 10 mL of Buffer L, bound proteins were eluted three times with 1 mL of elution buffer (Buffer L containing 150 mM imidazole, pH 8.0) and then dialyzed overnight in 1 L of starting buffer (10 mM Tris-HCl, pH 7.0, 50 mM NaCl). After replacing with new buffer, dialysis was continued for a further 20 h. The dialyzed sample was filtered and applied to a HiTrap Q column (GE Healthcare) and fractionated by linear gradient with end buffer (10 mM Tris-HCl, 1 M NaCl) using an AKTAprime plus (GE Healthcare) at 4°C. Proteins in each fraction were analyzed by 10–20% SDS-polyacrylamide gel electrophoresis and fractions exclusively containing a protein with the expected size of His-RNAP2 Ser2ph- mintbody were pooled. His-tag was removed using an Enterokinase Cleavage Capture kit (Novagen), according to the manufacturer’s instructions. Diluted recombinant enterokinase (10 µL) and 2 µL of 1 M CaCl2 were added to 1 mL of purified protein sample. After overnight incubation at room temperature, enterokinase was removed with EKapture™ Agarose. The concentration of the purified RNAP2 Ser2ph-mintbody was determined using Bradford protein assay (Bio-Rad) with bovine serum albumin (BSA) as the standard.

### ELISA

Microtiter ELISA plates were coated with 1 µg/mL BSA conjugated with RNAP2 CTD repeat peptides containing phosphorylated amino acids (Mab Institute Inc.; Table S1) overnight at 4°C, and were washed three times with PBS. Each well was incubated with 100 µL of Blocking one P (Nacalai Tesque) for 20 min at room temperature, washed with PBS, and then incubated with 50 µL of a 1:3 dilution series of purified RNAP2 Ser2ph-mintbody and IgG antibodies specific for Ser2ph (CMA602 and 42B3) and Ser5ph (CMA603) in PBS overnight. RNAP2 Ser2ph- mintbody and antibodies were diluted from 2.7 µg/mL and 300 ng/mL, respectively. Microtiter plates were washed three times with PBS, incubated with horseradish peroxidase (HRP)- conjugated anti-GFP (MBL; 1:2,000) and HRP-conjugated anti-mouse IgG (Jackson ImmunoResearch; 1:10,000) for RNAP2 Ser2ph-mintbody and IgG, respectively, for 90 min at room temperature. After washing four times with PBS, 100 µL of o-phenylenediamine solution (freshly prepared by dissolving one tablet containing 13 mg o-phenylenediamine-2HCl into 50 mL of 0.1 M sodium citrate buffer, pH 5.0, and 15 µL 30% hydrogen peroxide) was added. After incubating at room temperature until a yellow color was observed, the absorbance was measured at 490 nm, with a reference at 600 nm, using a Varioskan (Thermo Fisher Scientific).

### Live imaging of RNAP2 Ser2ph-mintbody and inhibitor treatments

Live-cell imaging of HeLa cells expressing RNAP2 Ser2ph-mintbody was performed using a high-resolution spinning disk confocal microscope featuring microlens-associated pinholes and a 3.2× tube lens to theoretically improve the optical resolution 1.4-fold (Olympus; IXplore SpinSR) equipped with a UPlanApo 60× OHR (NA 1.5) and a UPlanApoN 60× OSC2 (NA 1.4) objective lens for single- and multi-color fluorescence imaging, respectively, using a 488 nm laser (OBIS 488 LS; Coherent; 100 mW), a 561 nm laser (OBIS 561 LS; Coherent; 100 mW), an sCMOS (Hamamatsu Photonics, ORCA Flash 4), and a heated stage (Tokai Hit; 37°C, 5% CO2), under the operation of cellSens Dimension 3.1 software (Olympus).

To measure the number and size of RNAP2 Ser2ph-mintbody foci (Fig. 2, A–C), z-stack images were acquired using a 488 nm laser (100% transmission; 300 ms) at 0.2 μm intervals. Using ImageJ Fiji 1.52d software, line profiles of single foci were drawn and fit using a Gaussian distribution; Y = a + b*exp(-(x – c)^2^/(2d^2^)), to calculate the full width at half maximum (FWHM). To measure the number of foci per nucleus, foci were detected using NIS Elements (Nikon) using the spot detection function (binary > spot detection > bright spots with a typical diameter of 7 px) in all z-sections. The contrast in the spot detection function was adjusted to maximally detect nuclear spots without cytoplasmic spots. As one focus can be detected in three consecutive sections at the maximum, the total number of detected foci in the whole nucleus was divided by three.

For live-cell imaging of HeLa cells that stably expressed both RNA2 Ser2ph-minbody and H2B-Halo during mitosis (Fig. 2, D-F), cells were incubated in 50 nM HaloTag TMR ligand (Promega) for 30 min and the medium was replaced with FluoroBrite (Thermo Fisher Scientific) with supplements. Images were collected sequentially with a 488 nm laser (100% transmission; 300 ms) and a 561 nm laser (1% transmission; 300 ms) every 1 min.

For inhibitor treatments, after 10 images were acquired using a 488 nm laser (10% transmission; 200 ms) without intervals, flavopiridol (Sigma-Aldrich; at a final concentration of 1 µM), triptolide (Tocris; at a final concentration of 5 µM) or the vehicle (DMSO, 1:10,000 dilution) was added, and 10 images without intervals were acquired every 30 min for 2.5 h. The area of the RNAP2 Ser2ph-mintbody foci (Fig. 3) was measured using ImageJ Fiji software. Ten consecutive images were averaged, and the cytoplasmic average intensity was subtracted as the background. Auto-thresholding using the “Moments” mode was then applied to select the foci. When the cytoplasmic foci were selected, the “Remove Outliers” function with a radius of 0.2 was applied to remove the noise, and auto-thresholding was applied again. These procedures were repeated until cytoplasmic noises disappeared. The total area of the selected foci was calculated using the “Analyze Particles” function.

### Florescence recovery after photobleaching (FRAP)

HeLa cells stably expressing RNAP2 Ser2ph-mintbody were grown on a 35 mm dish and FRAP was performed in inhibitors that were added 2–4 h before the experiment. A dish was set on a heated stage (Tokai Hit) at 37°C under 5% CO2 (Tokken) on a point-scan confocal microscope (FV1000; Olympus) with a UPlan Apo 60× OSC (NA 1.4) lens under the operation of built-in ASW ver. 4.2 software (Olympus). Overall, 250 confocal images were collected every 47.8 ms (24 × 128 px; 2 μs pixel dwell time; zoom ×8.0; pinhole 800 μm; 0.2% transmission for a 488 nm line from a 20 mW multi-Ar+ ion laser), a 2 µm diameter circle was bleached using a 488 nm laser line with 100% transmission for 55 ms, and a further 50 images were collected using the original settings. The intensity of the bleached, nuclear reference, and background areas was measured using ASW ver. 4.2 (Olympus). After subtracting the background from the bleached and nuclear reference areas, the relative fluorescence in the bleached area was obtained by two step normalization. The intensity in the bleached area at each time point was obtained by dividing by that in the nuclear reference area, and the resulting relative intensity was then normalized using the average before bleaching.

### Expression and imaging of HaloTag-tagged proteins

To construct expression vectors for HaloTag-tagged proteins (CDK9, CDK12, and LEO1), total RNA was prepared from HeLa cells using TRIzol (Thermo Fisher Scientific) according to the manufacturer’s instructions, and cDNA was synthesized using the SuperScript III First-Strand Synthesis System for RT-PCR (Thermo Fisher Scientific). To the total RNA (2.3 µg in 1 µL), 7 µL of nuclease-free water, 1 µL of 50 µM oligo (dT)20, and 4 µL of 2.5 mM dNTP were added, and the mixture was incubated at 65°C for 5 min and chilled on ice for 1 min. After 4 µL of 5× First-Strand buffer, 1 µL of 0.1 M DTT, 1 µL of RNase OUT recombinant RNase inhibitor, and 1 µL of SuperScript III RT were added, the mixture was incubated for 60 min at 50°C and then for 15 min at 70°C, before the addition of 1 µL of RNase H (2 U/µL) and incubation for 15 min at 37°C. cDNA was purified using a PCR purification kit (Qiagen) and the concentration was measured using NanoDrop (Thermo Fisher Scientific). To amplify specific protein sequences, PCR was performed in a reaction mixture containing 5 µL of 5× Q5 buffer, 2 µL of 2.5 mM dNTPs, 0.6 µL each of forward and reverse primers (Table S1), 1 µL of cDNA, 15.55 µL of water, and 0.25 µL of Q5 DNA Polymerase (New England Biolabs), with the following PCR cycle: 1 min at 98°C, 35 cycles of 10 s at 98°C, 5 s at 72°C, 30 s at 72°C, and 2 min at 72°C. The PCR products were cloned into the HaloTag vector based on PB533 using an in-fusion system (TaKaRa). The expression vectors of H2B-Halo and Halo-PCNA were constructed by inserting their coding sequence (Kanda et al., 1998; Leonhardt et al., 2000) into the same HaloTag PB533 vector. The expression vectors of BRD4-Halo and p300-Halo were Kazusa clones (Promega). The expression vector of SF2(SRSF1/ASF)-mRFP has been described previously (Yomoda et al., 2008).

HeLa cells grown on glass bottom dishes were transfected with the expression vectors of HaloTag-tagged proteins and, the next day, cells were incubated in 50 nM HaloTag TMR Ligand (Promega) for 30 min, before washing with FluoroBrite (Thermo Fisher Scientific) containing supplements. Fluorescence images of sfGFP and TMR or mRFP1 were sequentially acquired using an IXplore SpinSR (Olympus) with a 488 nm laser (50% transmission; 300 ms) and a 561 nm laser (50% transmission for RPB3, BRD4, p300, SRSF1, PCNA, and H2B; 100% for CDK9, CDK12, and LEO1; 200 ms).

ImageJ Fiji software was used for colocalization analyses (Fig. 5). To draw a line profile, a 4 px-thick line was drawn and expressed as the relative intensity to the average nuclear intensity after subtracting cytoplasmic intensity. CCF was calculated using Just Another Colocalisation Plugin for ImageJ software. The Gaussian filter (radius 2.0 pixels) was applied to diminish the pseudo-correlation caused by pixel-to-pixel sensitivity variation, and the average fluorescence intensity of the nucleoplasm was subtracted from the image for thresholding. The CCF was measured with the x-shift set to 100 px (3.3 μm) without rotation and with rotation by 90°, 180°, and 270°, and the resulting four graphs were averaged and plotted in the range of −50 ≤ x ≤ 50 px. Pearson’s correlation of the nucleus was obtained using NIS Elements (Nikon).

### Single-molecule analysis

HeLa cells stably expressing RNAP2 Ser2ph-mintbody were transfected with expression vectors for HaloTag-tagged proteins (RPB3, BRD4, CDK9, p300, H2B, and PCNA) and were stained with 24 pM HaloTag TMR ligand for 30 min just before live-cell imaging. The medium was then replaced with FluoroBrite containing supplements. Single-molecule imaging was performed on a custom-build inverted microscope (IX83; Olympus) equipped with a 488 nm laser (OBIS 488 LS; Coherent; 100 mW), a 561 nm laser (OBIS 561 LS; Coherent; 100 mW), 100× objective lens (PlanApo NA 1.45 TIRFM; Olympus), and an intermediate magnification lens (2×; Olympus) (Tokunaga et al., 2008; Lim et al., 2018). HILO illuminations of both laser lines were achieved using a CellTIRF system (Olympus). Images were recorded at 33.33 ms per frame for 500 frames using two EMCCD cameras (C9100-13; Hamamatsu Photonics) and AQACOSMOS software (Hamamatsu Photonics). The spatial shift between the two camera images was corrected using the images of a 10 μm square lattice with ImageConverter software (Olympus). Cells were maintained at 37°C during observation using a stage top incubator and objective heater (Tokai Hit). Then, 15 cells were observed for each sample, and two independent experiments were performed.

Single-molecule tracking was carried out using modified u-track MATLAB code (Jaqaman et al., 2008). Briefly, single-molecule positions were detected using the detectSubResFeatures2D function in u-track and were precisely determined by two-dimensional Gaussian fitting. The localizations were linked using the trackCloseGapsKalmanSparse function with a maximum step length of 720 nm. Neither Gap closing nor Kalman filter in the u-track were used. The trajectories that lasted at least six steps (200 ms) were used to calculate MSD (Ito et al., 2017). The first six steps of MSD were fit with MSD = *4Dt* to determine the diffusion coefficient (*D*) of an individual trajectory; as the period of analysis was very short (200 ms), the simple diffusion model was used. The maximum threshold of the diffusion coefficient *D*thr for bound molecules was determined from the distribution of log10(*D*). A stacked histogram from all samples was fitted to a double Gaussian function, and the mean minus two standard deviations of the fast fraction was used as *D*thr. Similar values for *D*thr (0.065 and 0.063 μm^2^/s) were obtained from two independent replicate experiments.

The relative intensity of RNAP2 Ser2ph-mintbody was obtained from a 500-frame (16.67 s) averaged image that was recorded simultaneously with single-molecule imaging. Non-uniform lighting of the average image was minimized by a 2 μm high-pass image filter. The range of the intensity from minus to plus two standard deviations from the mean was normalized to that from 0 to 1. The normalized intensity of the mintbody at each pixel point on the trajectory of HaloTag-tagged protein was calculated. Since each trajectory traverses various mintbody intensity regions, the average of the top two intensities of the points on each trajectory was used as the relative RNAP2 Ser2ph-mintbody intensity (*I*rel_Ser2ph) of the trajectory. Based on the images, trajectories with the top 25% and bottom 25% *I*rel_Ser2ph were predominantly located inside and outside RNAP2 Ser2ph foci, respectively. The difference between the *D*s of bound molecules (≤ *D*thr) inside (the top 25% *I*rel_Ser2ph) and outside (the bottom 25% *I*rel_Ser2ph) RNAP2 Ser2ph foci was analyzed using the Mann-Whitney U test.

### Tracking RNAP2 Ser2ph-mintbody and Cy3-DNA foci

Cells expressing RNAP2 Ser2ph-mintbody were bead-loaded with 0.1 mM Cy3-dUTP (Perkin-Elmer) (Manders et al., 1999; Sato et al., 2018) and cultured for two days. On the day of imaging, the medium was replaced with FluoroBrite (Thermo Fisher Scientific) containing supplements. Fluorescence images of sfGFP and Cy3 images were sequentially acquired at 551 ms/frame using an IXplore SpinSR (Olympus) with a 488 nm laser (50% transmission; 300 ms) and a 561 nm laser (10% transmission; 200 ms).

For tracking foci, RNAP2 Ser2ph and Cy3 foci were detected with NIS Elements (Nikon) using binary > spot detection > bright spots with the following parameters: typical diameter pixels 10 (RNAP2 Ser2ph) or 12 (Cy3); contrast 4.94 (RNAP2 Ser2ph) or 2.52 (Cy3); maximum gap size 3; and delete tracks <30 frames. Mean-square displacements (MSDs) up to 20 steps (11 s) were used for fitting using MSD = 4*Dt^α^*, where *D*, is the diffusion coefficient; *t* is the elapsed time; and α is the anomalous exponent (0 < α < 2).

### Data availability

The nucleotide sequence data of 42B3(R78K/A80T/M95V)-scFv are available in several public databases (DDBJ/EMBL/GenBank) under the accession number LC628457.

## Acknowledgements

We thank M.C. Cardoso for PCNA construct, H. Ochiai for valuable comments, members of Kimura lab for constructive discussion, and the Biomaterials Analysis Division, Open Facility Center, Tokyo Institute of Technology for DNA sequencing analysis.

## Author contributions

Conceptualization, H.K.; Investigation, S.U., Y. I., Y.S., and H.K.; Formal analysis, S.U. and Y.I.; Methodology, Y.I. and Y.S.; Resources, Y.S., T.H., Y.O., and M.T.; Writing–original draft, S.U., Y.I., and H.K.; Writing–review and editing, Y.S., T.H., Y.O., and M.T.; Funding acquisition, Y.I., Y.S., Y.O., T.H., M.T., and H.K.; Supervision, H.K.

## Summary of Supplementary Materials

Table S1. Primers used in this study.

Figure S1. Screening of RNAP2 Ser2ph-specific scFv-sfGFP.

Figure S2. Development of RNAP2 Ser2ph-mintbody.

Figure S3. Localization of various proteins respect to RNAP2 Ser2ph-mintbody.

Figure S4. Scatter plots of the diffusion coefficient of HaloTag-tagged proteins and the relative RNA2 Ser2ph-mintbody intensity.

Movie 1. RNAP2 Ser2ph-mintbody and Halo-H2B during prophase to prometaphase.

Movie 2. RNAP2 Ser2ph-mintbody and Halo-H2B during telophase to G1 phase.

## Movie legends

**Movie 1. RNAP2 Ser2ph-mintbody and Halo-H2B during prophase to prometaphase.** Single confocal sections of HeLa cells that stably express RNAP2 Ser2ph-mintbody (green) and Halo-H2B (magenta) were acquired every 1 min. RNAP2 Ser2ph-mintbody foci that were observed in the early prophase disappeared as the prophase progressed. RNAP2 Ser2ph- mintbody diffused into the cytoplasm after the nuclear membrane broke down.

**Movie 2. RNAP2 Ser2ph-mintbody and Halo-H2B during telophase to G1 phase.** Single confocal sections of HeLa cells that stably expressed RNAP2 Ser2ph-mintbody (green) and Halo-H2B (magenta) were acquired every 1 min. RNAP2 Ser2ph-mintbody was excluded from condensed chromosomes during the telophase, but became concentrated in small foci as the nucleus was formed. The number of foci increased as the G1 phase progressed.

---

**Table S1.**
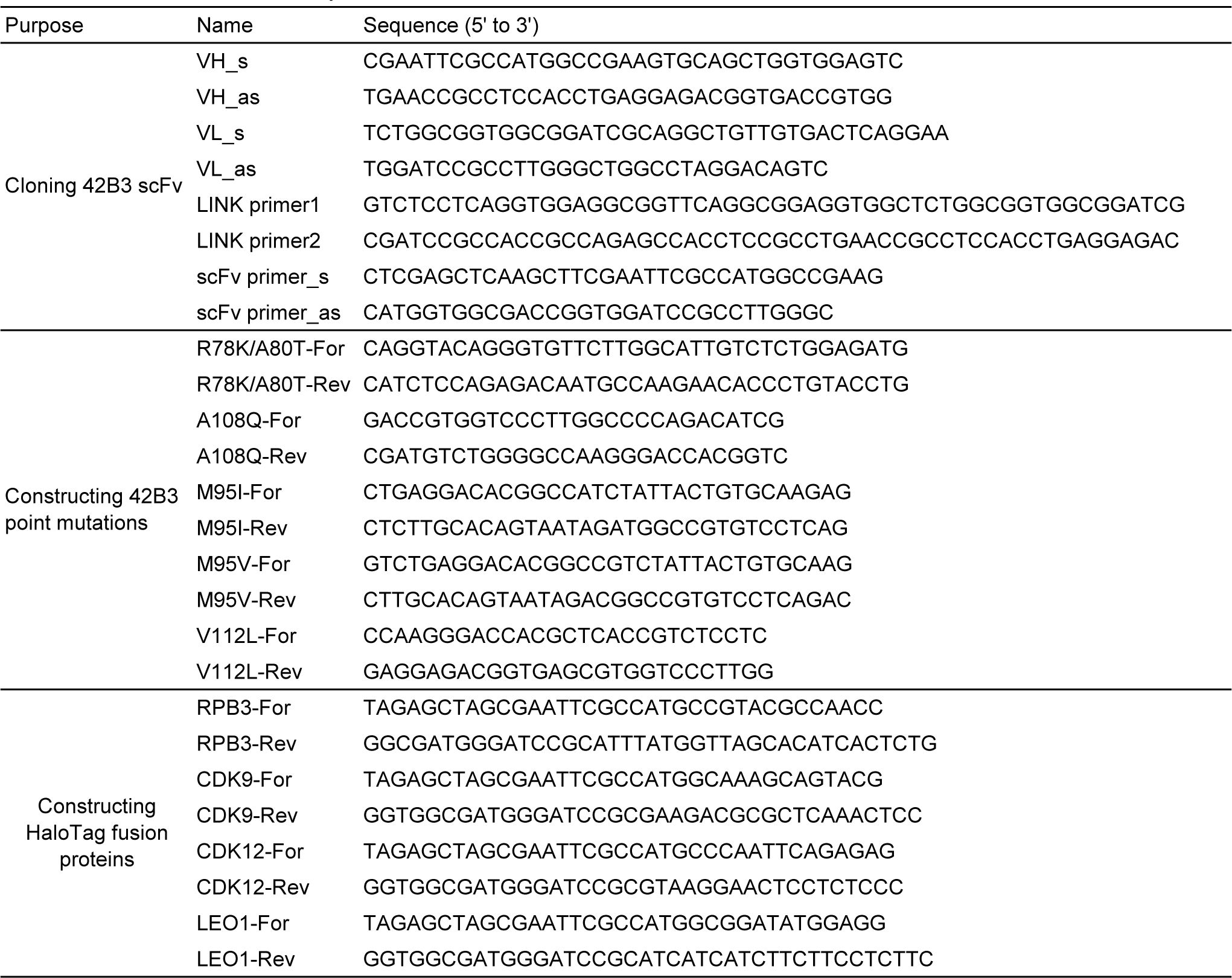
Primers used in this study.

**Figure S1.**
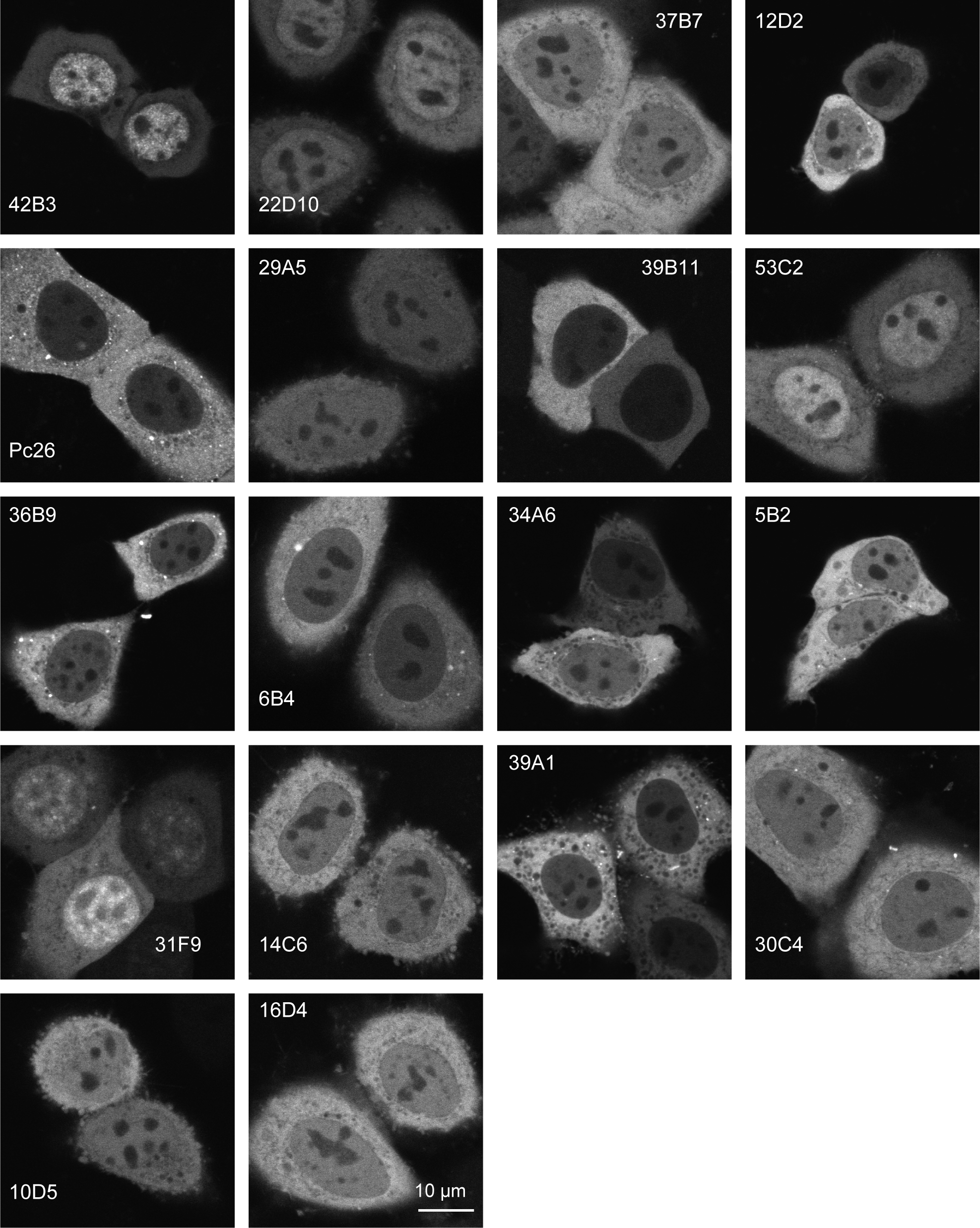
Screening of RNAP2 Ser2ph-specific scFv-sfGFP. Single-chain variable fragments tagged with sfGFP (scFv-sfGFP) were transiently expressed in HeLa cells and the fluorescence patterns were imaged using a confocal microscope. Single confocal sections are shown. Among the 18 different clones tested, 42B3 was most accumulated in the nucleus, where the target RNAP2 Ser2ph is present. Scale bar, 10 µm.

**Figure S2.**
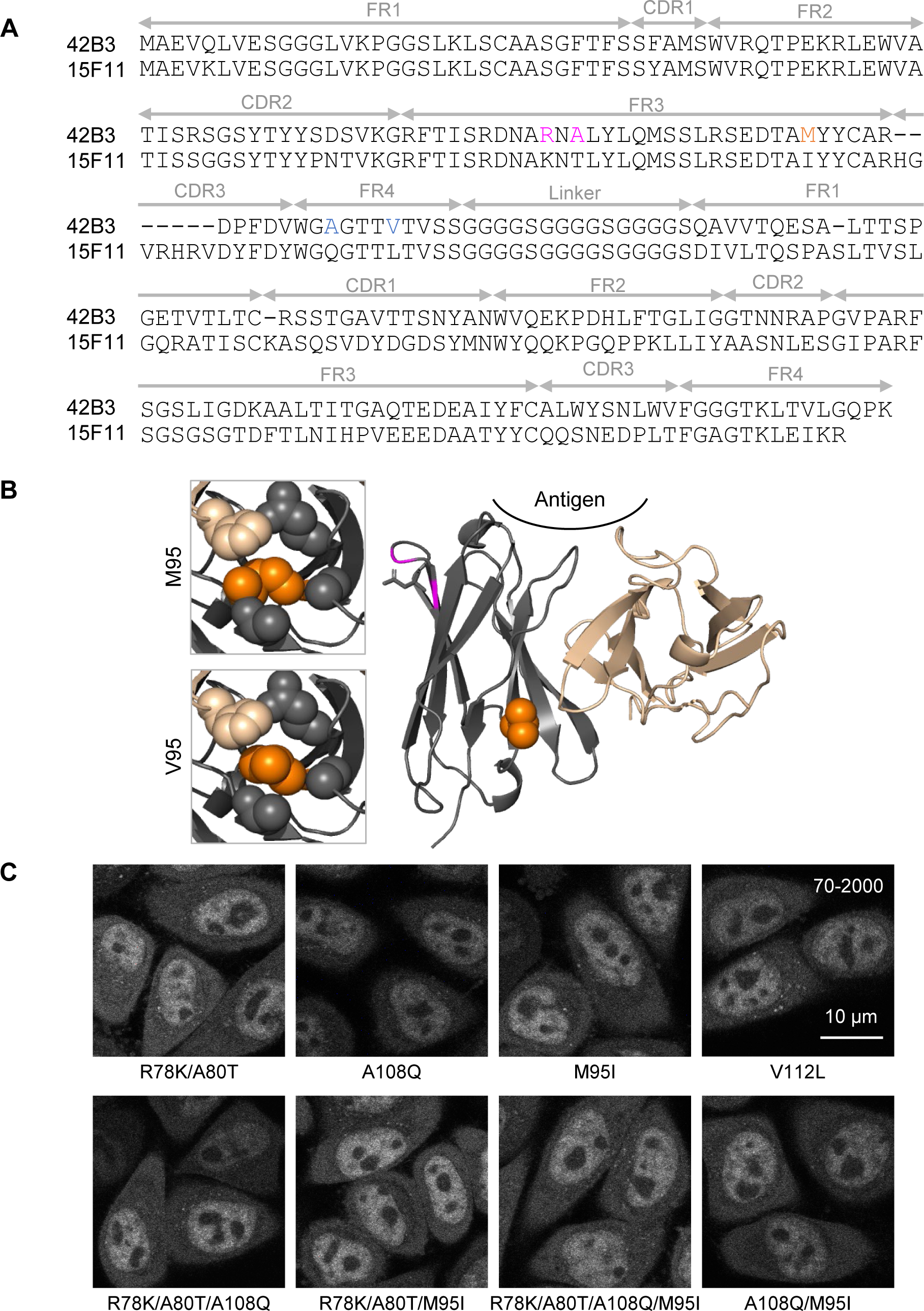
Development of RNAP2 Ser2ph-mintbody. (A) Amino acid sequence of RNAP2 Ser2ph (42B3) and H4K20me1 (15F11) scFvs. Framework regions (FRs), complementarity determining regions (CDRs), and the linker region are indicated. Mutated sites in 42B3 are indicated in colors. The final construct named RNAP2 Ser2ph-mintbody contains R78K, A80T, and M95V substitutions. (B) Comparison of model structures of 42B3 and 42B3(R78K/A80T/M95V). The models were generated using 15F11 (PDB: 5B3N). The M95V mutation appears to strengthen the hydrophobic core more than the original 42B3. (C) Example images of mutant scFv-mCherry that were stably expressed in HeLa cells. The image acquisition setting and contrast adjustment (70-2000) are the same. Scale bar, 10 µm.

**Figure S3.**
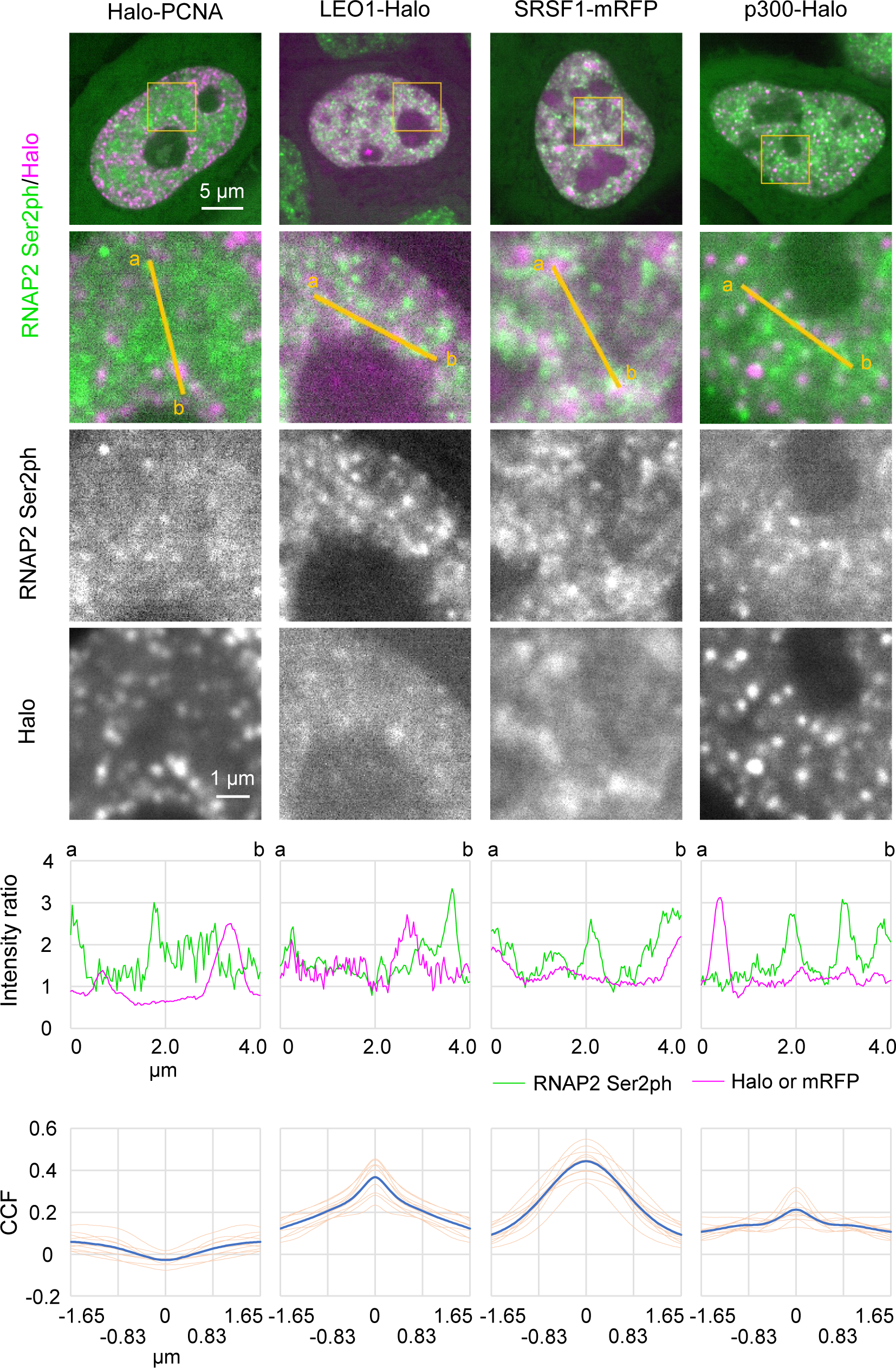
Localization of various proteins respect to RNAP2 Ser2ph-mintbody. HeLa cells stably expressing RNAP2 Ser2ph-mintbody were transfected with expression vectors for HaloTag-tagged proteins (p300, LEO1, and PCNA) or SRSF1-mRFP, and stained with 50 nM HaloTag Lignad-TMR for 30 min before live-cell imaging using a high-resolution confocal system. Averaged images of 10 consecutive frames (551 ms/frame) are shown with magnified views of areas indicated in orange (middle three row) and normalized profiles of 4 µm lines (fourth). Cross-correlation functions of 10 cells are shown at the bottom (blue, the average; orange, individual cells). Scale bar, 5 µm.

**Figure S4.**
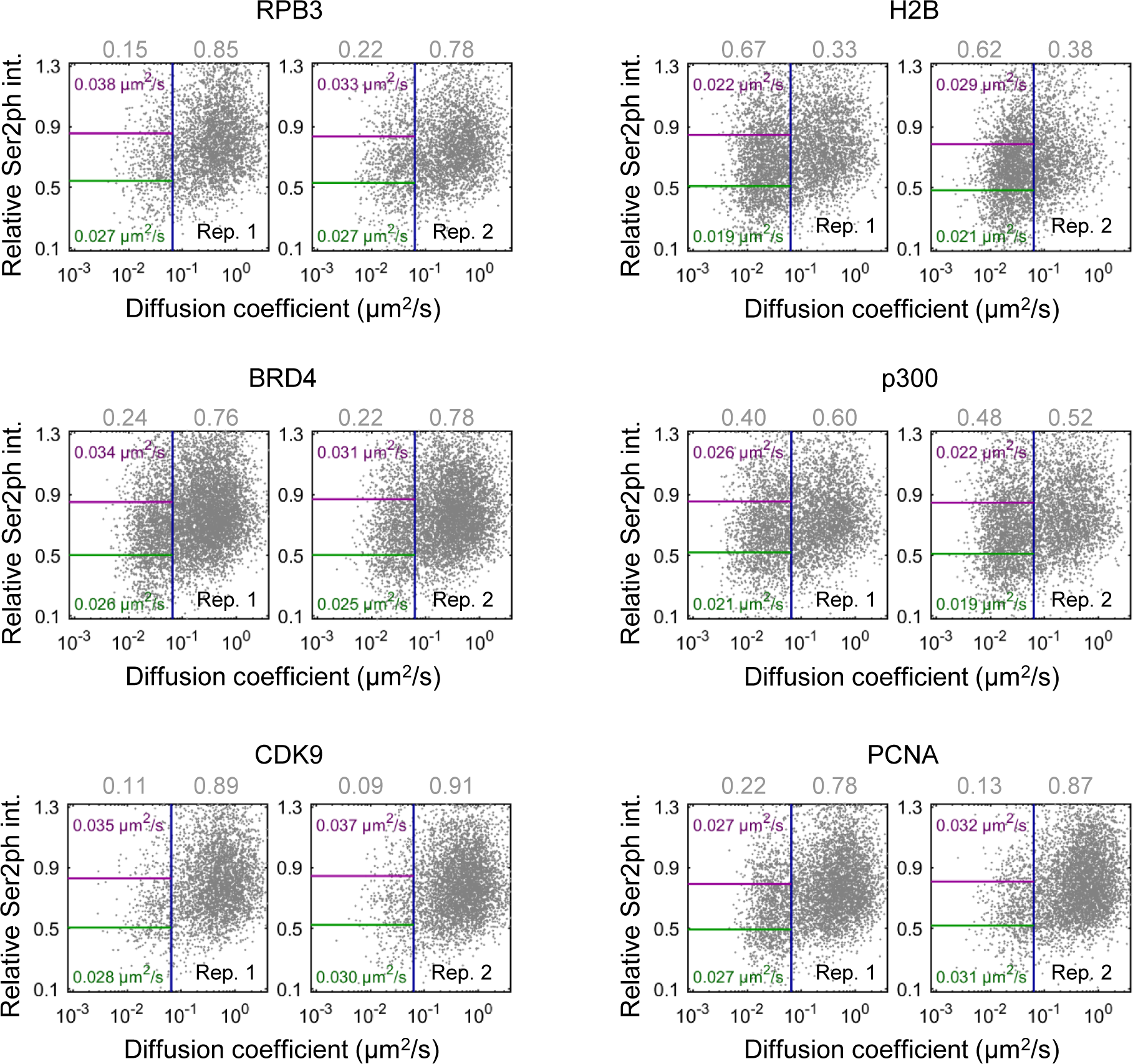
Diffusion coefficient of HaloTag-fusion proteins and RNAP2 Ser2ph-mintbody intensity. Scatter plots of the diffusion coefficient *D* and the relative RNAP2 Ser2ph-mintbody intensity. The diffusion coefficients (*D*s) of proteins were obtained using single-molecule trajectories of HaloTag-tagged proteins (RPB3, BRD4, CDK9, p300, H2B, and PCNA) stained with HaloTag TMR Ligand recorded at 33.33 ms/frame, which were superimposed upon highpass-filtered RNAP2 Ser2ph-mintbody images in living Hela cells (Fig. 6). Scatter plots of *D* and the relative RNAP2 Ser2ph-mintbody intensity *I*_rel_Ser2ph_ from two independent replicate experiments are shown. Each dot represents a single trajectory (molecule). The ratio of the bound (*D* ≤ *D*_thr_) and mobile (*D*_thr_ < *D*) fractions is shown in gray above the plots. The average *D*s of bound molecules of the top and bottom 25% *I*_rel_Ser2ph_ are shown in magenta and green, respectively. A blue vertical line indicates *D*_thr_.

